# Exposure to pyrethroid insecticides modulates immunity of *Anopheles* against *Plasmodium falciparum*

**DOI:** 10.1101/2025.07.30.667449

**Authors:** Patrick Hörner, Leonard Heereman, Christoph Wenz, David Steeb, Lorenz Decius, Juliane Hartke, Alisa Botsch, Julia Bettina Mäurer, Ralf Krumkamp, Victoria Anne Ingham

## Abstract

Transmission of malaria parasites relies on the blood-feeding behaviour of female *Anopheles* mosquitoes and can be prevented by vector control methods. Particularly, the usage of pyrethroid-containing insecticide-treated nets (ITNs) remains a cornerstone of malaria prevention. Whilst ITNs are primarily designed to induce mosquito mortality, their impact on parasite development, particularly in the context of pyrethroid resistance, is poorly understood. Our study reveals that sub-lethal pyrethroid exposure triggers systemic increases in reactive oxygen and nitrogen species reducing transmission efficacy of *Plasmodium falciparum*. Production of reactive nitrogen species in granulocytes induces their proliferation and primes mosquito immunity through activation of a non-canonical immune deficiency (IMD) pathway, leading to increased nitration around the midgut epithelium and subsequent ookinete destruction. These findings highlight an overlooked secondary mode of action for pyrethroids and reinforces the importance of sustained pyrethroid usage in ITNs, even in the face of high-levels of pyrethroid resistance in mosquitoes.

## Introduction

Organisms undergo continuous evolution to better shield themselves from abiotic and biotic stressors. This is exemplified in the insect vectors of disease which must adapt to overcome insecticide use and defend themselves against the pathogens by which they are infected. Adaptations to such environmental and organismal pressures substantially changes the underlying cellular biology of the insect. The deadliest vector-borne disease is malaria caused by the parasite *Plasmodium falciparum*, infecting over 250 million people and resulting in nearly 600 000 deaths in 2023, mostly in African children under 5 ^1^. Transmission control efforts have revolved heavily around the deployment of insecticide-treated bednets (ITNs) that contain insecticides of the pyrethroid class aiming to kill the *Anopheles* mosquito vectors. The molecular mechanisms of resistance employed by anophelines to these insecticides are complex, including physiological and metabolic changes ^2^, such as increased expression of the detoxifying enzymes cytochrome P450s (P450s) and glutathione S-transferases (GSTs) ^3,4^. In turn, changes in basal metabolic activity and respiration have been shown in pyrethroid resistant mosquitoes and other insect pest species ^5,6^. Furthermore, these key cellular processes are perturbed upon exposure to pyrethroid insecticides in multiple insects, indicating off-target responses to these neurotoxic insecticides ^7,8^. Increases in respiration are likely occurring through alterations to mitochondrial oxidative phosphorylation ^5,8^, which could lead to oxidative stress through extensive generation of superoxide anions (O2^.^-) ^9,10^; thus, presumptively disrupting redox homeostasis within the mosquito. Disruption to redox homeostasis leads to oxidative damage through production of deleterious amounts of reactive oxygen species (ROS) and reactive nitrogen species (RNS) ^11–13^. ROS and RNS are oxidants with short half-lives and play crucial roles in cellular signaling by reversibly modifying biomolecules ^14–16^. In the context of mosquito biology, ROS can profoundly affect mosquito fitness ^17–19^ insecticide susceptibility ^20^, immunity against bacteria ^21^, arboviral infection ^22^ and *Plasmodium* development ^21,23,24^, whilst research on RNS has been mostly limited beyond their coincidental function in immune defense ^23,25,26^.

After an infectious blood meal, the *Plasmodium* ookinete must transverse the gut epithelium of the *Anopheles* mosquito to establish infection; this occurs around 18-30 hours post-blood meal ^27–29^. At this stage, *Plasmodium* will encounter the mosquito innate immune system, including haemocytes, the immune cells of the mosquito circulating in the haemolymph ^30^. Initiation of an effective early phase immune response is reliant on provisions of high local levels of hydrogen peroxide (H_2_O_2_) from NADPH oxidase 5 (NOX5) and nitric oxide (NO) catalyzed by nitric oxide synthase (NOS) from L-arginine ^31–33^. Subsequent formation of highly toxic intermediates induces epithelial nitration of midgut cells and sparks a multifaceted complement response ^34^. The complement response begins with the recruitment of phagocytic haemocytes (granulocytes) to the basal lamina and their release of haemocyte-derived microvesicles (HdMv’s) during caspase-mediated apoptosis ^34^ and ultimately culminates in parasite lysis, likely after deposition of mosquito-protein on the parasite surface ^35^. In addition to lysis, the parasite can be encapsulated by melanin, which can be augmented through continuous suppression of midgut catalase during invasion, which sustains high local H_2_O_2_ levels ^21,36,37^.

The immune deficiency pathway (IMD), and to a lesser extend the c-Jun N-terminal kinase (JNK) pathway, drive the production of key effector molecules, including thioester-containing protein 1 (TEP1), in tissues and granulocytes targeting *Plasmodium*, whereas JNK predominantly mediates the epithelial nitration response in the mosquito midgut ^38–40^. This epithelial nitration cascade and ensuing complement response are deficient in defending against *P. falciparum* in insecticide-susceptible mosquitoes from the same geographical region, attributed to disruption of JNK signaling, leading to immune evasion ^41–43^. However, it remains unclear whether the altered basal metabolism in insecticide-resistant mosquitoes or their acute response to insecticide exposure can impact this critical anti-parasitic defence. Given the high likelihood of infected, pyrethroid-resistant *Anopheles* encountering insecticides multiple times during a single infection cycle, understanding the cellular changes in the context of parasite development is crucial. Previous research has demonstrated that pyrethroid exposure leads to increased ROS in different biological systems ^13,44^ and that resistant *Anopheles* have evolved to effectively manage the ensuing endogenously elevated H_2_O_2_ levels ^20^. The relationship between vector competence and perturbations in redox state are further complicated by changes to the midgut microbiota caused by ROS ^21^, which have been shown to play a key role in *Plasmodium* infection ^45–47^. Despite the critical importance of pyrethroid insecticides in malaria control, the link between immunity and resistance has remained understudied due to the challenges of controlled infections with well characterised, insecticide resistant mosquitoes. Indeed, only two studies have directly explored the impact of pyrethroid exposure on parasite development, one in natural infections showing reduced levels of *P. falciparum* ^48^ whilst a resistant *Anopheles* line created through back-crossing into a susceptible background showed no impact of exposure on infection; however, the resistant mosquitoes sustained higher levels of infection ^49^.

Here, we demonstrate that insecticide resistance mechanistically influences vector competence in a major African *Anopheles* species following exposure. We show that exposure to the widely used pyrethroid insecticide permethrin significantly alters both ROS and RNS levels, thereby disrupting redox homeostasis in critical tissues involved in the infection response to *Plasmodium*. Further, we find changes to the composition of circulating mosquito immune cells and the critical immune factor TEP1 after insecticide exposure, leading to reduced oocyst numbers, indicating that insecticides have a direct impact on immunity. We then show that artificial increase of RNS levels leads to decreased vectorial capacity in pyrethroid resistant and susceptible mosquitoes through increased nitration and ookinete lysis. Accompanying RNAseq reveals that increased RNS impacts known anti-*Plasmodium* immunity genes in resistant mosquitoes and results in proliferation of haemocytes involved in the complement cascade, potentially through a non-canonical IMD -controlled pathway. Finally, we indicate that the microbiome is unlikely to be driving this phenotype. In conclusion, our findings offer new insights into the intricate interplay between insecticide resistance, redox homeostasis and immunity in malaria vectors. These findings have critical implications on the continued use of pyrethroid insecticides for malaria control and highlight the potential for targeting the redox system in transmission control strategies.

## Results

### Insecticide resistance impacts ROS-/RNS levels in Anopheles mosquitoes

To explore whether pyrethroid resistance leads to increased ROS or RNS in *Anopheles* mosquitoes, we compared constitutive transcript expression of the RNS-producing enzyme, nitric oxide synthase (NOS), the RNS-limiting enzyme, arginase, and the major ROS detoxifying enzymes, catalase (CAT) and three superoxide dismutases (SODs) in two pyrethroid resistant populations (PY-R) and one pyrethroid susceptible (PY-S) population of the *Anopheles gambiae* species complex. We found significantly higher *NOS* mRNA expression in both resistant populations: Tiefora (*An. coluzzii* - 5.9-fold (*p_T-TEST_=0.0012*)); Tiassalé (*An. gambiae* – 3.4-fold (*p_T-TEST_=0.0131*)) (Figure 1A) compared to the susceptible Kisumu (*An. gambiae*) population, whilst arginase expression was mildly elevated in the Tiefora PY-R population (Figure S1A). *CAT* also showed a significantly increased expression in both resistant populations: Tiefora - 1.9-fold (*p_T-TEST_=0.0092*), Tiassalé (*An. gambiae* s.l.) – 1.6-fold (*p_T-TEST_=0.0121*) (Figure 1B), whilst none of the *SOD* isoenzymes were overexpressed (Figure S1A). Due to the high levels of *NOS* expression, we then determined differences in protein nitration levels, a proxy for RNS activity, in PY-R and PY-S mosquitoes using dot blots with an anti-nitrotyrosine antibody. In line with the expression data, we saw a slight but non-significant increase in total RNS in both PY-R populations (Figure S1B). As there were no observable differences in the two resistant populations, we continued with only one PY-R mosquito population (Tiefora, *An. coluzzii*). We compared ROS and RNS levels in PY-R and PY-S mosquitoes in tissues related to *Plasmodium* development: the midgut and salivary glands. In contrast to whole mosquitoes, midguts of PY-R mosquitoes showed 1.5-fold lower levels of nitrotyrosine (*p_T-TEST_=0.0324*; Figure S1C) and also significantly 2.4-fold lower ROS levels (*p_T-TEST_<1e-4*; Figure S1D), whilst salivary glands showed no change (Figure S1E-F) compared to the susceptible counterpart. These data demonstrate that insecticide resistance causes changes to the oxidative state of mosquitoes, particularly in the redox sensitive midgut.

**Figure 1.**
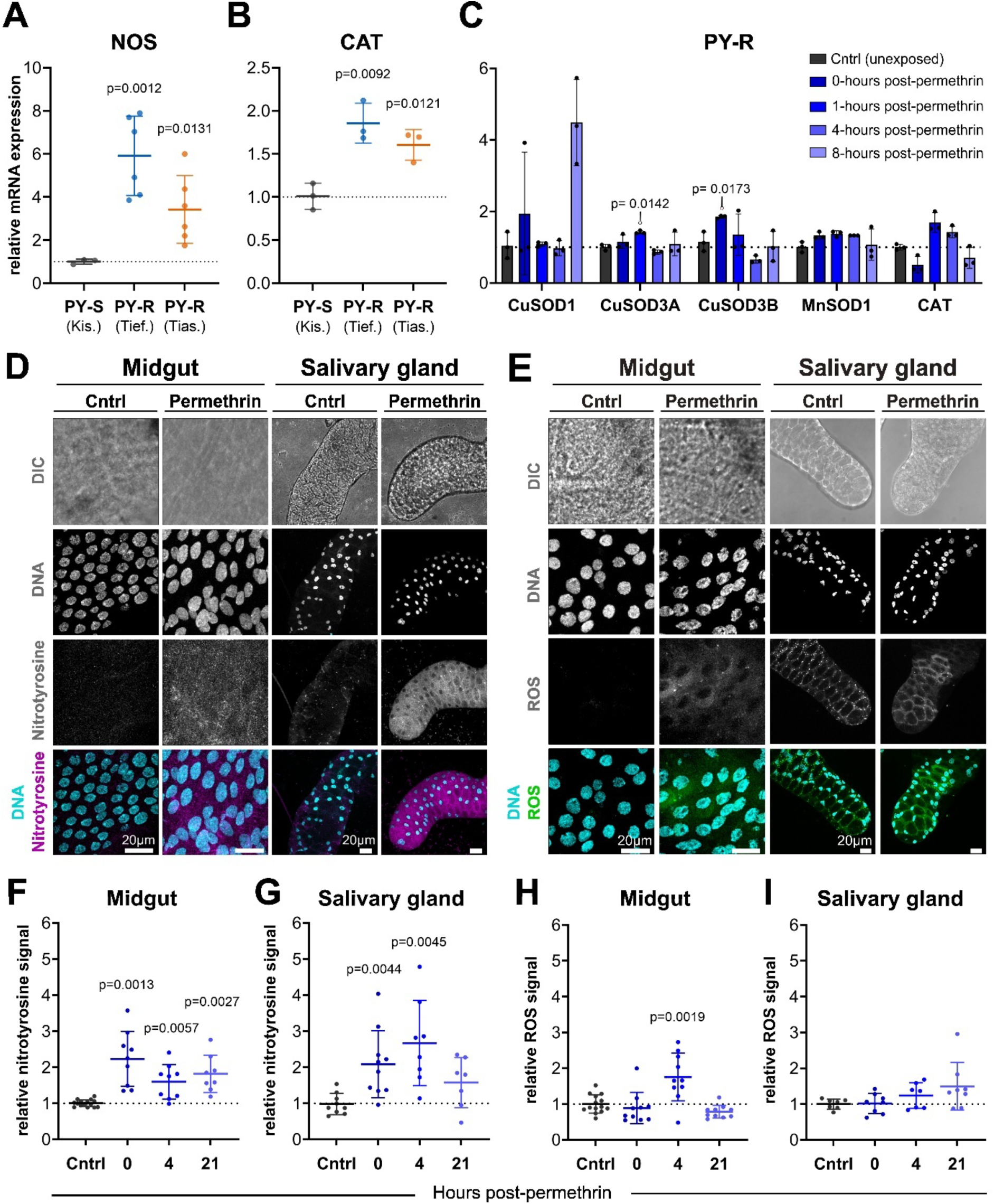
Permethrin triggers RNS and ROS generation in Plasmodium-related tissues. (A and B) Relative mRNA expression levels of NOS (A) and CAT (B) (y-axis) in two pyrethroid-resistant (PY-R) populations (Tiassalé – Tias., Tiefora – Tief.) compared to a susceptible (PY-S) population (Kisumu – Kis.) (x-axis). Significance: t-test with Bonferroni correction (cut-off: 0.025). (C) Relative mRNA expression (y-axis) of ROS and RNS related transcripts at multiple timepoints (0 to 8 hours) following 0.75% permethrin exposure (1 hour) of PY-R Tiefora (x-axis). Significance: one-way ANOVA with Dunnett’s test. Error bars in A-C show standard deviation, each point is one biological replicate of seven pooled mosquitoes. (D and E) Confocal images showing protein nitration (D) and ROS (E) in midguts and salivary gland lobes of PY-R Tiefora following exposure to 0.01% permethrin (30 min tarsal) in acetone or to acetone-only (Cntrl). Representative images show peak nitration or ROS levels after exposure. Gray = Brightfield, cyan = DNA staining (Hoechst), magenta = nitrated proteins (anti-nitrotyrosine antibody), green = ROS (CellROX™ Deep Red). Scale bar = 20 µm. (F and G) Quantitative analysis of the nitrotyrosine fluorescence signal (y-axis) in midguts (F) and salivary glands (G) of exposed PY-R mosquitoes at three timepoints (0, 4 and 21 hours) after insecticide contact relative to Cntrl (x-axis). (H and I) Quantitative analysis of the ROS fluorescence signal (y-axis) in dissected midguts (H) and salivary glands (I) of exposed PY-R mosquitoes at three timepoints (0, 4 and 21 hours) after insecticide contact relative to Cntrl (x-axis). Error bars in F-I show standard deviation; each point is one midgut or salivary gland. Significance: t-test with Bonferroni correction (cut-off: 0.0167).

### Pyrethroid exposure boosts ROS-/RNS levels in Plasmodium-related tissues

To determine whether pyrethroid insecticides trigger ROS and/or RNS generation, we explored the effects of a sub-lethal pyrethroid insecticide dose on ROS/RNS levels in the PY-R population, from immediately after exposure up to 21-24 hours, the approximate time of ookinete traversal. We exposed mosquitoes to the diagnostic dose of permethrin ^50^ and found significant increases in mRNA of the ROS detoxifying enzymes *SOD3B* immediately (1.9-fold; *p_ANOVA_=0.0173*) and *SOD3A* at 1 hour (1.4-fold; *p_ANOVA_=0.0142*) post-exposure, whilst *CAT* showed an obvious upward trend between 1- and 4 hours post-exposure and *SOD1* at 8 hours post-exposure (Figure 1C). *NOS,* whose expression remains incompletely understood in insects, appears to regulate itself via distinct isoforms that induce or suppress its expression through transcriptional and post-translational mechanisms ^51–53^, showed no increase in mRNA expression up to 24 hours post-exposure (Figure S1G). However, we detected an increase in total full-length NOS protein of 1.6-fold immediately, a peak 6.2-fold increase at 4 hours and slowly reverting back to 3.7-fold 21 hours post treatment by western blot (Figure S1H), when we used an LC_30_ dose of permethrin (Figure S1I). With the same insecticide dose, we also quantified nitric damage in the midgut and salivary glands and saw a clear increase in protein nitration in both tissues (1.5x - 3x that of the control), starting immediately after the 30 min exposure (0-hour timepoint) up to at least 21 hours post-exposure (Figure 1D, 1F, 1G). Midguts had significantly increased nitric damage across all time points, whilst the salivary glands recovered by 21-hours. We also used a generic ROS dye and detected a specific ROS burst (1.75-fold increase) in midguts 4-hours post-permethrin exposure, whilst salivary glands showed a slight upwards trend from 4-hours onwards but were not significantly affected (Figure 1E, 1H, 1I). Taken together, pyrethroid exposure clearly shifts the redox balance towards higher oxidative stress, including a sustained RNS increase in *Plasmodium*-relevant tissues.

### Pyrethroid exposure decreases vector competence by modulating Anopheles immunity via ROS/RNS

Phagocytic haemocytes, the granulocytes, are important for activation of the complement system in the early phase of the anti-plasmodial immune response ^34^. Unexpectedly, we noticed that exposure to the LC_30_ permethrin dose caused a highly significant increase in apparent blebbing of haemocytes during the imaging of mosquito immune cells at all investigated timepoints (Figure S1J-K), indicating that the pyrethroid has a direct impact on their cellular integrity/composition. To determine whether insecticide exposure could also impact haemocyte composition, we determined the proportion of the three major haemocyte populations (prohaemocytes, oenocytoids, granulocytes) by imaging after extracting the cells from the haemolymph ^54,55^. In addition to visual discrimination by granularity and size, we also confirmed classification as granulocytes by using the lipophilic cell labelling dye Vybrant CM-Dil, which is preferentially taken up by granulocyte populations (Figure S1L) ^34,56^. We found significant increases in granulocyte proportions at all timepoints (Figure 2A), with a peak increase from 10.8% in the control to 31.9% at 21 hours post-exposure (*pΞ* ^2^*<1e-4*). For this reason, we focused on this crucial cell type in the following experiments. To investigate oxidative and nitrosative effects on these cells, we again quantified protein nitration and ROS levels ^57^. We observed no differences in neither RNS nor ROS basal levels between PY-R and PY-S mosquitoes (Figure S1M-N); however, increased protein nitration after contact with permethrin was evident across all time points (Figure 2B, 2D; *p_MW_=2e-4 (0 hours), p_MW_=0.0124 (4 hours), p_MW_<1e-4 (21 hours*)). In contrast to the midgut where an immediate ROS burst was observed, here increased ROS (3.9-fold increase; *p_MW_<1e-4*) was evident at 21 hours post-exposure (Figure 2C, 2D). The increased proportion and nitration of granulocytes also led us to investigate whether the TEP1-dependent complement response is affected, as this parasite lysis mediating protein is primarily produced in granulocyte clusters and the fat body, where immune cells are formed ^58^. Indeed, total full-length TEP1 protein (∼165 kDa) and the processed/cleaved version TEP1-cut (∼80 kDa) were both increased after permethrin exposure at all timepoints ^35^, with protein levels peaking immediately after the 30 min exposure (0-hour timepoint) at approximately 2.5-fold (Figure 2E). We also detected a third unknown band around 70 kDa uniquely at the 4-hour time point and another around 60 kDa at all timepoints which both could be caused by autocatalysis of the protein.

**Figure 2.**
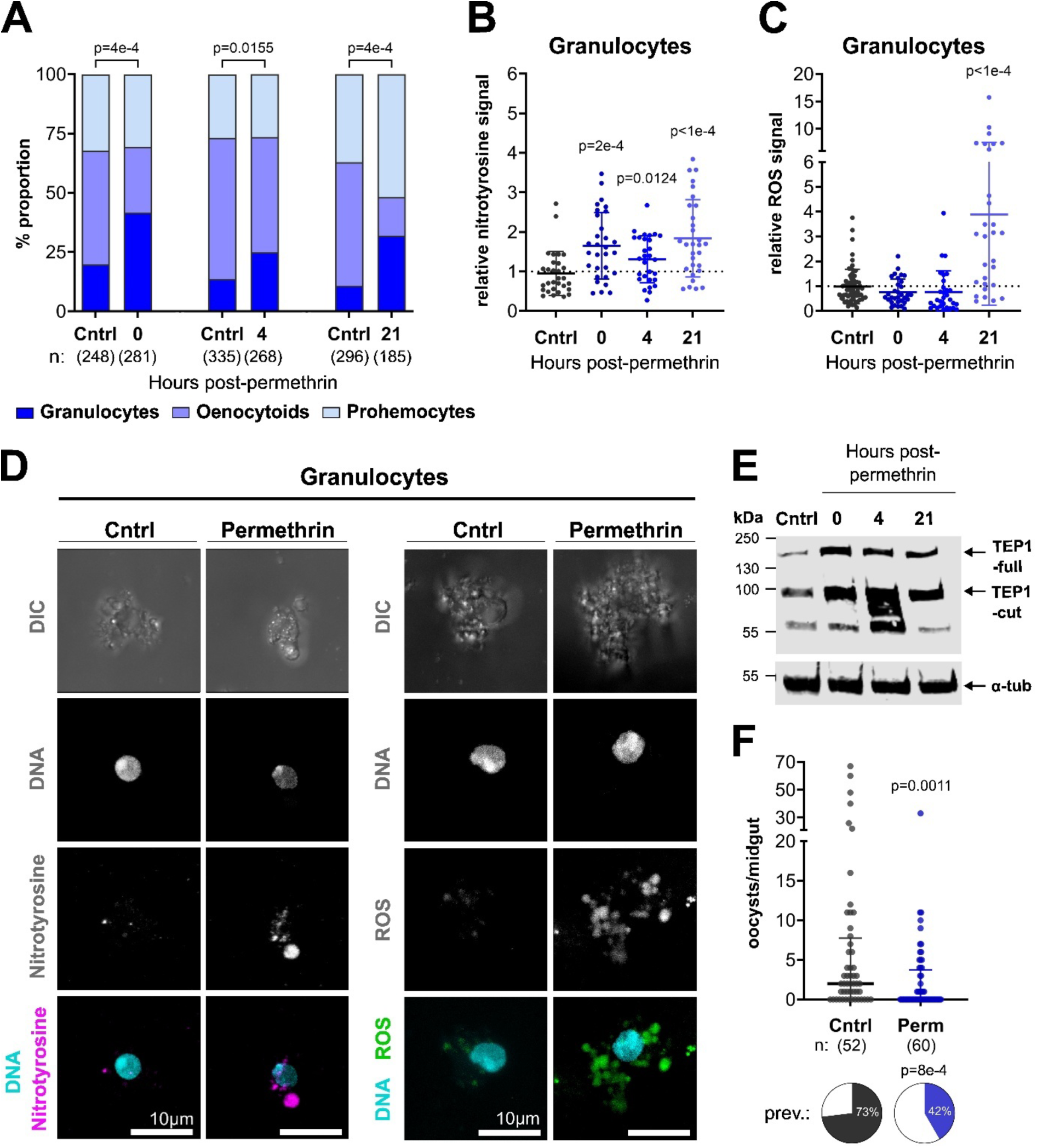
Permethrin exposure reduces vector competence by modulating immune cell populations and the mosquito complement system. (A) Percentage of immune cell populations (granulocytes, oenocytoids, prohemocytes) determined via confocal imaging (y-axis) at three timepoints (0-, 4- and 21 hours) after tarsal contact of PY-R to 0.01% permethrin in acetone (30 min) or acetone-only control (Cntrl) for the respective timepoint (x-axis). n = total number of counted cells per group. Significance: chi-square test. (B and C) Quantitative analysis of the nitrotyrosine fluorescence signal (B) and ROS signal (C) (y-axis) in extracted granulocytes of exposed PY-R mosquitoes at three timepoints (0-, 4- and 21 hours) after insecticide contact (x-axis). Error bars show standard deviation, each point is one granulocyte. Significance: Mann Whitney test with Bonferroni correction (cut-off: 0.0167). (D) Confocal images showing protein nitration and ROS in extracted granulocytes of PY-R mosquitoes after tarsal exposure to 0.01% permethrin in acetone (30 min) or to acetone-only (Cntrl). Representative images show peak nitration and ROS levels after exposure. Gray = Brightfield, cyan = DNA staining (Hoechst), magenta = nitrated proteins (anti-nitrotyrosine antibody), green = ROS (CellROX™ Deep Red). Scale bar = 10 µm. (E) Western blot showing total TEP1 protein (TEP1-full = full length protein, TEP1-cut = cleaved/activated version) and α-tubulin (internal loading control) at three timepoints (0-, 4- and 21 hours) after 0.01% permethrin contact of pools of five PY-R mosquitoes for 30 min (x-axis). (F) Oocysts per midgut (dot plot) and infection prevalence (prev.; pie chart) at 5 days post Plasmodium falciparum infection in PY-R after tarsal exposure to 0.01% permethrin (Perm) for 7.5 min in acetone or to acetone-only (Cntrl). n = number of midguts per treatment group. Each point represents oocyst number in one midgut from three individual infection replicates. Significance: Mann-Whitney test (oocysts per midgut) and chi-square test (infection prevalence).

As permethrin exposure causes clear changes to the redox imbalance in *Plasmodium*-related tissues and changes to immune response, we then determined whether insecticide exposure directly impacted *Plasmodium* development. Critically, a single tarsal exposure to permethrin followed by infection of *P. falciparum* caused a significant decrease in oocyst intensity and prevalence (Figure 2F), whilst oocyst size was not affected (Figure S1O). Taken together, these results point to large effects of sub-lethal pyrethroid exposure on the mosquito immune system, possibly impacting parasite development before oocyst establishment.

### Targeted RNS increases modulate mosquito immunity via granulocyte nitration

To establish if the observed changes in the mosquito immune system and parasite development after permethrin exposure are directly linked to oxidative-/nitrosative stress, we investigated the impacts of direct RNS and ROS manipulation on these phenotypes.

To explore nitrosative stress, we utilized a highly conserved process: the oxidation of L-arginine (L-Arg) by NOS to L-citrulline producing nitric oxide (NO), the most basic RNS ^59^. To this end, we administered L-Arg to mosquitoes over their entire life time via addition to cotton wool pads, and first established the highest dose of L-Arg (0.2%) that does not affect mosquito longevity (Figure S2A). This dose was used for all further experiments. A dot blot of whole mosquitoes revealed that L-Arg treatment leads to a significant increase of nitrotyrosines (1.4-fold; *p_T-TEST_=0.0492;* Figure S2B), demonstrating that L-Arg treatment increases nitrosative stress. We then checked for protein nitration in all parasite-related tissues and found a highly significant (*p_MW_<1e-4*), almost 2-fold increase in nitrotyrosine signal specifically in immune cells (Figure 3A and 3B), mimicking the response to permethrin (Figure 2B). Interestingly, we observed no increase in nitrosative damage in the midgut or salivary glands (Figure S2C-D).

**Figure 3.**
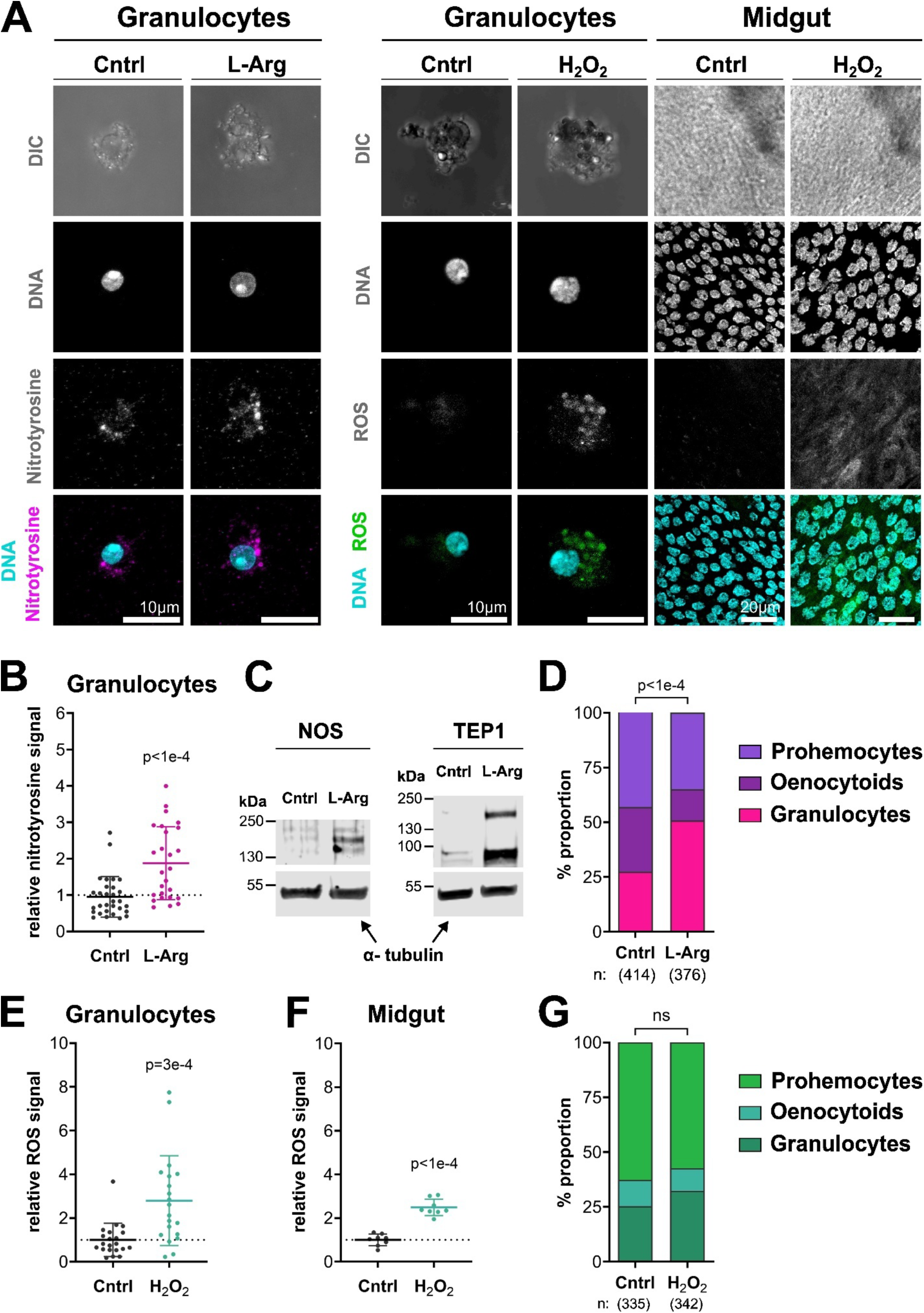
L-arginine induces RNS in granulocytes and increases their proportion while hydrogen peroxide affects ROS but not immune cell populations. (A) Representative confocal images showing protein nitration in granulocytes and ROS in granulocytes and midguts of PY-R control-(Cntrl), L-arginine (L-Arg) or hydrogen peroxide (H_2_O_2_) fed. Gray = Brightfield, cyan = DNA staining (Hoechst), magenta = nitrated proteins (anti-nitrotyrosine antibody), green = ROS (CellROX™ Deep Red). Scale bar = 10 µm (granulocytes), 20 µm (midguts). (B) Quantitative analysis of the nitrotyrosine fluorescence signal (y-axis) in extracted granulocytes of PY-R receiving Cntrl or L-Arg diet (x-axis). Error bars show standard deviation; each point is one granulocyte. Significance: Mann-Whitney test. (C) Western blots showing NOS and TEP1 protein (*α*-tubulin = internal loading control) of Cntrl or L-Arg fed PY-R. (D) Percentage of immune cell populations (granulocytes, oenocytoids, prohemocytes) determined via confocal imaging (y-axis) in Cntrl or L-Arg fed PY-R (x-axis). n = total number of counted cells per group. Significance: chi-square test. (E and F) Quantitative analysis of the ROS fluorescence signal (y-axis) in extracted granulocytes (E) and midguts (F) of Cntrl or H_2_O_2_ fed PY-R Tiefora mosquitoes (x-axis). Error bars show standard deviation; each point is one granulocyte or midgut. Significance: Mann-Whitney test in (E) and t-test in (F). (G) Percentage of immune cell populations (granulocytes, oenocytoids, prohemocytes) determined via confocal imaging (y-axis) in Cntrl or H_2_O_2_ treated PY-R (x-axis). n = total number of counted cells per group. Significance: chi-square test (ns = not significant).

As pyrethroid exposure altered haemocyte composition and led to an increase in TEP1 levels, we determined whether L-Arg similarly impacted immunity and the immune cells. We saw a 3-fold increase in total *NOS* mRNA expression (Figure S2E), 3.9-fold in total protein (Figure 3C) and large increases of 7.8-fold in full-length TEP1 and 5-fold in TEP1-cut (Figure 3C). We again observed a highly significant shift in the proportion of circulating granulocytes in the whole immune cell population, similar to the response at 21-hours post permethrin exposure, with an increase in granuloycyte proportion from 27.5 to 50.8% upon L-Arg treatment (*pχ* ^2^*<1e-4*) (Figure 3D). Taken together, these data show that L-Arg treatment mimics the immune changes seen after pyrethroid exposure.

### Increasing ROS levels impacts Plasmodium-related tissues and granulocytes

Although it is very difficult to replicate the timepoint-specific ROS bursts we saw in tissues after permethrin exposure, we used the same feeding approach for ROS manipulation to determine their antiparasitic effects. First, we determined a suitable dose for H_2_O_2_ supplementation (0.15%) (Figure S2F) and subsequently quantified ROS levels via imaging of parasite-related tissues. We found a highly significant increase in ROS levels in both the granulocytes and midguts (Figure 3A, 3E, 3F), with a 2.8-fold and 2.5-fold increase in the mean ratio respectively (*p_MW_=3e-4; p_T-TEST_<1e-4*), whilst no change was seen in the salivary glands (Figure S2G). We again classified haemocyte proportions but found no significant shift in the main populations by H_2_O_2_ (Figure 3G), indicating that this response seems to be mediated by RNS but not ROS levels in immune cells.

### RNS changes in immune cells inhibits oocyst formation through ookinete lysis

We next explored how the observed secondary impacts of pyrethroid exposure, namely RNS and ROS induction, impacted vector competence for the human malaria parasite *P. falciparum* in both PY-R and PY-S mosquitoes. In contrast to the very effective nitric oxide signaling response against the rodent model malaria parasite *P. berghei* ^25^, *P. falciparum* can evade this immune response during midgut invasion by ookinetes ^60–63^. To determine whether increased RNS levels impact *P. falciparum*, we scored infection in the mosquito midgut after L-Arg treatment at an early (5 days post-infectious blood mean (dpIBM) and later time point (10 dpIBM). We discerned highly significant decreases in oocyst intensity at both early and late timepoints in both PY-R (*p_MW_=0.0055; p_MW_<1e-4*) and PY-S (*p_MW_=0.015; p_MW_=5e-4)* (Figure 4A) mosquitoes. Combining the data from both timepoints, we found that L-Arg treated PY-R and PY-S mosquitoes were significantly less likely to establish infection by 1.7-(risk ratio: 95% confidence interval [CI] = 1.32 to 2.27) and 1.2-(risk ratio: 95% confidence interval [CI] = 1.02 to 1.33) times, respectively (Figure 4B). L-Arg had no impact on oocyst size in either population (Figure S3A-B), indicating at a specific impact against the early phases of the infection before oocyst establishment occurs. In line with the reduced infection rate, L-Arg provision also led to significantly fewer salivary gland sporozoites in both PY-R (*p_MW_=0.0021*) and PY-S (Figure 4C; *p_MW_ =0.0088*) mosquitoes, and less infectious mosquitoes (PY-R - 1.9-fold; PY-S – 1.3-fold).

**Figure 4.**
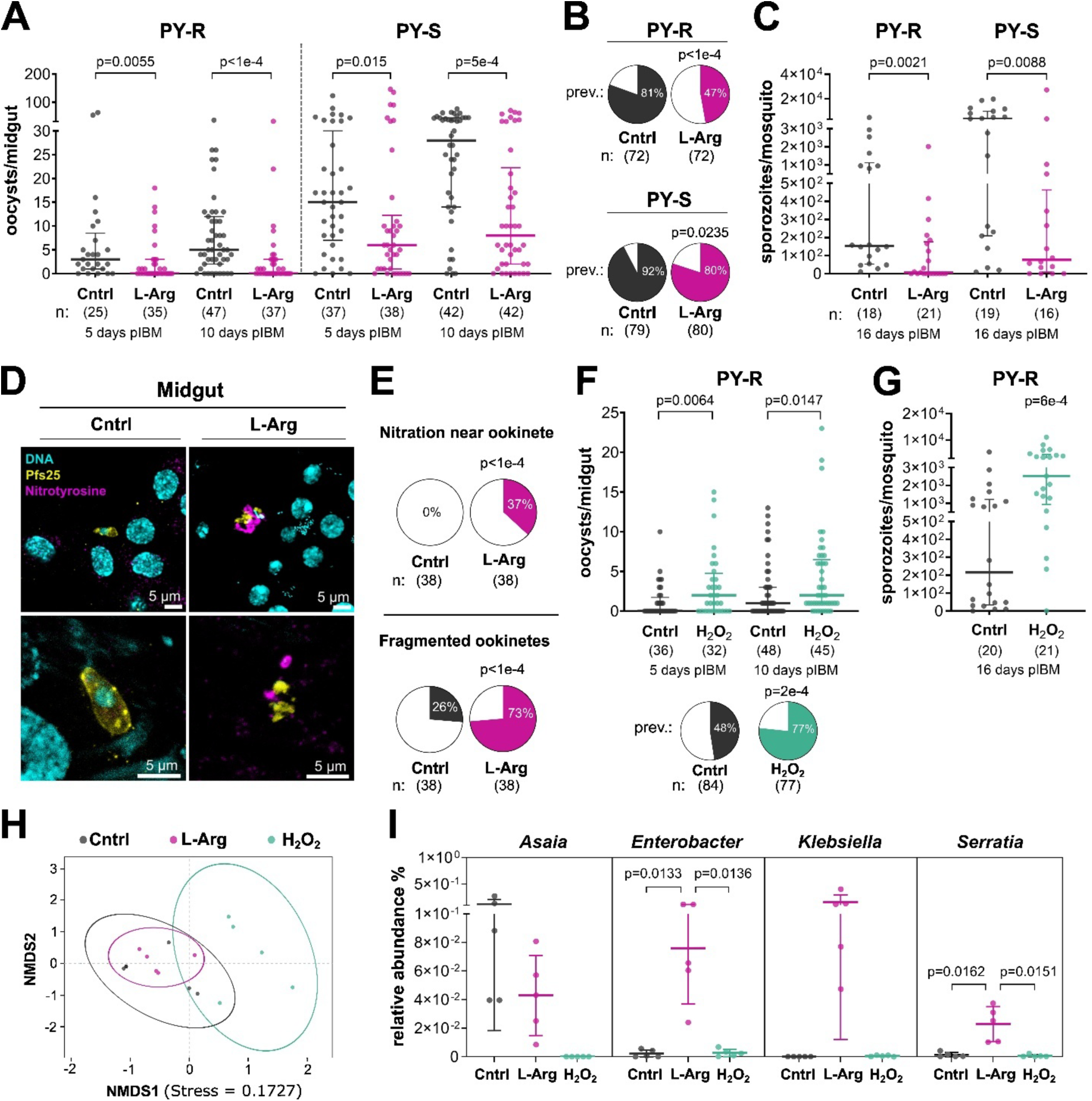
L-arginine inhibits P. falciparum development in insecticide resistant and susceptible mosquitoes through ookinete lysis while H_2_O_2_ promotes development. (A) Oocysts per midgut in PY-R and PY-S (y-axis) at 5 days post-infectious blood meal (pIBM) and 10 days pIBM control (Cntrl) or L-arginine (L-Arg) fed. n = number of midguts per timepoint and treatment. Each point is oocyst count of one midgut of at least three individual infectious feeds. Significance: Mann-Whitney test. (B) Infection prevalence (prev.) in PY-R and PY-S combining both oocyst timepoints from (A). n = total number of midguts per treatment. Significance: chi-square test. (C) Salivary gland sporozoites per mosquito (y-axis) in Cntrl and L-Arg fed PY-R and PY-S at 16 days pIBM (x-axis). Each point is average of salivary gland sporozoites per mosquito of pools of 1-3 mosquitoes. Significance: Mann-Whitney test. (D) Confocal images showing ookinetes (yellow = anti-Pfs25 antibody), protein nitration (magenta = anti-nitrotyrosine antibody) and nuclei (Cyan = DNA) in midguts of Cntrl or L-Arg treated mosquitoes 26 hours post infection. Scale bar = 5 µm. (E) Percentages of ookinetes with clear nitration signal in proximity (upper) and fragmented ookinetes (lower) in Cntrl and L-Arg treated mosquitoes on the total number of imaged ookinetes (x-axis). n = total number of imaged ookinetes per treatment. Significance: chi-square test. (F) Oocysts per midgut (dot plot) and infection prevalence (pie chart) in PY-R (y-axis) at 5 days pIBM and 10 days pIBM in control (Cntrl) or hydrogen peroxide (H_2_O_2_) fed. n = number of midguts per timepoint and treatment. Each point is oocyst count of one midgut of at least three individual infectious feeds. Significance: Mann-Whitney test (oocysts per midgut) and chi-square test (infection prevalence). (G) Salivary gland sporozoites per mosquito (y-axis) in Cntrl or H_2_O_2_ fed PY-R at 16 days pIBM (x-axis). Each point is average of salivary gland sporozoites per mosquito of pools of 1-3 mosquitoes. Significance: Mann-Whitney test. (H) Non-metric multidimensional scaling plot of 16S bacterial analysis in Cntrl (black), L-Arg (pink) and H_2_O_2_ (green) treated PY-R mosquitoes 21 hours after a blood meal showing NMDS1 (x-axis) and NMDS2 (y-axis) for the calculated abundances. (I) Relative abundance on the whole microbiome of four different bacterial genera (Asaia, Enterobacter, Klebsiella, Serratia) in Cntrl. L-Arg, or H_2_O_2_ treated PY-R mosquitoes 21 hours after receiving a non-infectious blood meal. Each point is one biological replicate of pools of five mosquitoes. Significance: t-test with Bonferroni correction (cut-off: 0.0167)

As we did not observe any effect on oocyst growth including sporozoite formation and we detected no evidence of oocyst melanisation, we hypothesized that L-Arg affects an earlier stage of infection, the ookinete. To test this hypothesis, we infected mosquitoes with *P. falciparum* and stained the midguts with an ookinete-specific antibody and the nitrotyrosine antibody 24-26 hours pIBM. We observed a highly significant 2.8-fold increase in fragmented parasites compared to control ^64^, a proxy for lysis (Figure 4D-E; *pχ* ^2^*<1e-4*). Furthermore, we detected protein nitration in areas in direct proximity to the ookinete (Figure 4D-E) only in the L-Arg treated mosquitoes. Collectively, L-Arg-induced RNS elevation demonstrates direct ookinete-killing activity against *P. falciparum*.

### ROS induction promotes P. falciparum development

In contrast to L-Arg treatment, when PY-R mosquitoes were fed with H_2_O_2_ we saw increased early-(*p_MW_=0.0064*) and late-stage oocyst numbers (*p_MW_=0.0147*) after infection with *P. falciparum* (Figure 4F). Investigating infection prevalence also showed that H_2_O_2_ increased the likelihood of establishing an infection significantly by 1.6 times (Figure 4F; risk ratio: 95% confidence interval [CI] = 1.257 to 2.105). The average size of oocysts per midgut was not affected by the treatment (Figure S3C), indicating that there were no developmental defects during maturation. As a consequence of the increased infection rate, H_2_O_2_ increased transmission potential as sporozoite numbers were also significantly elevated (Figure 4G; *p_MW_=6e-4*). Previous research on rodent malaria parasites showed that silencing *catalase* in mosquitoes reduced the oocyst load in *Anopheles gambiae* ^21^, whilst other natural vector-parasite combinations in central America showed the opposite ^65^. Here, when we silenced *catalase* by RNAi (Figure S3D), there was no effect on oocyst numbers, prevalence nor size (Figure S3E-G). These results clearly suggest that a sustained, systemic elevation of ROS levels enhances vector competence but without impacting immune cell composition.

### ROS but not RNS cause large compositional shift of the microbiota

Previously changes in ROS levels have been linked to changes in the mosquito microbiota ^66^ and aseptic microbe-free conditions could directly impact *Plasmodium* infection ^47^, whilst the influence of RNS on the microbiome remains understudied. To determine if the changes in vectorial capacity could be due to changes in the microbiota, we performed 16S sequencing on L-Arg and H_2_O_2_ fed mosquitoes (Table S2). Whilst alpha diversity was not changed by either treatment (Figure S4A), analysis of beta diversity revealed large impacts on species abundance, shown in the non-metric multi-dimensional scaling (NMDS) analysis, after H_2_O_2_ feeding (Figure 4H). A Bray-Curtis plot that shows the distribution of the dissimilarity highlighted clear separation of the H_2_O_2_ group from the control group (Figure S4B) and significantly decreased diversity (*P_PERMANOVA_=0.001*). Interestingly, we found *Asaia spp.* that has been linked to impeding *Plasmodium* infections ^67,68^, completely eradicated by H_2_O_2_ (Figure 4I, Figure S4C); thus, we cannot rule out these changes impacting infection. Although L-Arg grouped closely with the control, there were three bacterial genera that showed a large increase in abundance after treatment, *Serratia, Enterobacter* and *Klebsiella* (Figure 4I, Figure S4C).

### Transcriptomic data shows wide-scale changes to major immune pathways after RNS treatment

To explore molecular processes involved in redox manipulation, parasite development and insecticide resistance, we performed RNA sequencing on PYR mosquitoes from different treatment groups (L-Arg, H_2_O_2_) and compared transcript expression to the respective non-supplemented controls. We used RNA extracted from *P. falciparum* infected mosquitoes around the time of ookinete invasion of the epithelium (21hrs pIBM), the developmental stage most impacted by ROS/RNS.

In L-Arg treated PY-R mosquitoes, differential gene expression analysis showed up-regulation of 2578 genes (24%) and downregulation of 2834 genes (26%) compared to the control (Table S3). GO term analysis revealed significant enrichment in oxidative phosphorylation (*p=4e-9*), cellular respiration (*p=1.03e-4*), structural component of the ribosome (*p=2.74e-15*) and, as expected, cellular nitrogen compound processes (*p=6.05e-6*). We next subset the transcript list to look at genes involved in immunity and found 146 immune-related genes significantly differentially expressed (81 up- and 65 down-regulated) and additionally 89 insecticide resistance-related genes (35 up- and 54 down-regulated) (Figure S5A-B; Table S4).

When exploring the transcripts related to immunity, we saw all major immune signaling pathways against *Plasmodium* were significantly down-regulated across multiple pathway hierarchies (Figure 5A). All members of the JAK/STAT pathway were down-regulated except the inhibitors *SOCS* and *PIAS*. Both the TOLL and JNK pathways were down-regulated from the point of signal initiation across all three hierarchical levels. The only exceptions were *PELLE* in the TOLL pathway and *MKK4* in the JNK pathway, which showed no change in expression. Notably, the negative regulators of both pathways, *PVR* and *PUC*, were also not differentially expressed. The most notable of the pathway changes occurs in the IMD pathway, where activation of multiple effector proteins is still evident, including antimicrobial peptides like *gambicin*, *cecropin1* and the major complement factors *TEP1*, *APL1C* and *LRIM1* ^35,69–72^. Furthermore, *IMD* itself is upregulated, as are the pathway activators *DIAP2* and *UEV1A*; however, the *REL1* transcription factor is down-regulated potentially indicating a non-canonical response ^73,74^. We also explored whether there was an enrichment in granulocyte markers of multiple previously identified granulocyte clusters (Table S5) ^58^, and found significant cluster membership, with the highest in haemocyte cluster 2 (65%), identified as granulocytes and the antimicrobial granulocyte cluster 6 (70%).

**Figure 5.**
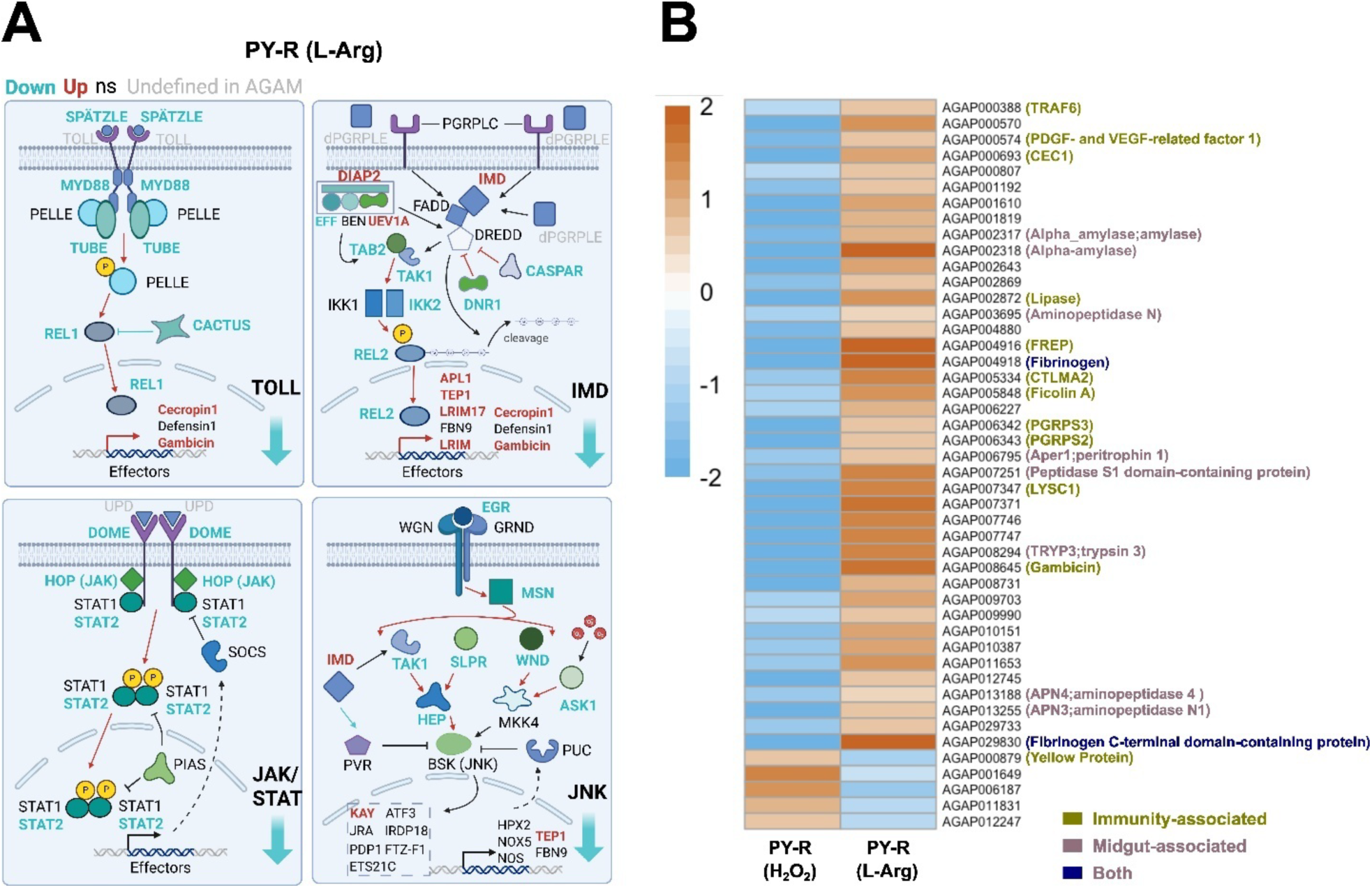
L-arginine induces expression of immunity effectors during ookinete invasion while hydrogen peroxide suppresses immune- and midgut-related genes. (A) RNA sequencing schematic showing up- and down-regulated genes of major immune signaling pathways against Plasmodium in PY-R L-Arg treatment group against Cntrl 21 hours post-infection. Blue = down-regulated genes, red = up-regulated genes, black = not significant and gray = undefined in Anopheles gambiae complex mosquitoes. (B) Log_2_ fold change for immunity- and midgut-associated genes differentially expressed in inverse direction 21 hours post-P. falciparum infection of PY-R treated with L-Arg or H_2_O_2_. Treatments were compared to respective Cntrl treated mosquitoes.

Compared to L-Arg-treated mosquitoes, transcriptome changes were much less pronounced in H_2_O_2_-treated PYR-R mosquitoes (Figure S5C-D, Table S6). In total, we only found 23 genes (0.2%) overexpressed compared to the control but 124 genes down-regulated (1.1%). Interestingly, GO term analysis of all down-regulated genes revealed defence response to other organism (*p=0.0027*), suggestive of a weakened immune response. This finding was supported by down-regulation of multiple immunity-related genes, including three CLIP-domain serine proteases (CLIPs) that are mostly involved in the insect melanisation response and in activation of the TOLL pathway ^75,76^.

When focusing on genes with opposite expression patterns within the H_2_O_2_ and L-Arg groups, we found multiple immunity-related and midgut-associated genes (Figure 5B; Table S7). Prominent examples were again antimicrobial peptides of the IMD- and TOLL pathways like *cecropin1* and *gambicin* but also *fibrinogen* and *fibrinogen-*related proteins (FREPs), that are linked to immunity against parasites ^77^, and a haemocyte recruitment factor (AGAP000574 - *PDGF-and VEGF-related factor 1*), indicating these transcripts could play a key role in the observed changes in vectorial capacity. Taken together, the transcriptomic changes induced by targeted RNS or ROS elevation during ookinete invasion reveal substantial shifts in genes and pathways associated with the early phase immunity.

### Modeling permethrin exposure and L-Arg uptake shows impacts on parasite transmission and human infection rates

So far, our results suggest that permethrin exposure or dietary L-Arg negatively affect vectorial capacity in laboratory settings. To quantify the potential impact of these treatments on malaria transmission in the field, we developed a compartmental SEIR (Susceptible-Exposed-Infected-Recovered) model with a human and a mosquito population, parameterised with field data from southern Burkina Faso, the origin of the *An. gambiae* PY-R (Tiefora) population used in this study (Figure S6A) ^78,79^. The model integrates experimentally derived effects of permethrin and L-Arg on PY-R mosquito fecundity, including egg output, hatch rate, and larval emergence, using solvent (acetone) and glycine as respective controls, and assumed no additional mortality from treatment (Figure S6B–D). To simulate the impact on transmission, we incorporated oocyst prevalence data from mosquitoes exposed to LC_30_ permethrin or fed 0.2% L-Arg (Figures 2F, 3B).

We first modelled increasing coverage of sub-lethal permethrin or L-Arg exposure across a single transmission season. Increasing permethrin coverage to 50–100% led to substantial declines in predicted human infections, particularly during peak transmission months (July–November) (Figure 6A), whereas L-Arg yielded more modest reductions (Figure 6B). Assuming full treatment coverage, both interventions shifted the population towards fewer infected and more susceptible individuals (Figure S6E–F). Next, we modelled seasonal variation by altering the baseline transmission rate, which reflects biting frequency and transmission efficiency ^79^. Under low transmission scenarios, representing dry season or lower endemicity, sub-lethal permethrin exposure was highly effective, whilst L-Arg again conferred milder benefits (Figure 6C–D; Figure S6G). Notably, permethrin retained efficacy even outside the peak season, underscoring the added benefit on reduced parasite development in areas with fluctuating vector densities. Finally, we extended the model to stratify the human population by age (Figure S6H), enabling predictions of intervention impact by demographic group (Figure 6E–F; Tables S8–9). Children under five, the most vulnerable group, showed the largest benefit. Permethrin treatment of 50% of mosquitoes reduced infections in this age group by 12.8%, rising to 31.2% with 100% exposure. L-Arg supplementation yielded reductions of 8.7% and 22.4%, respectively, for the same treatment coverages.

**Figure 6.**
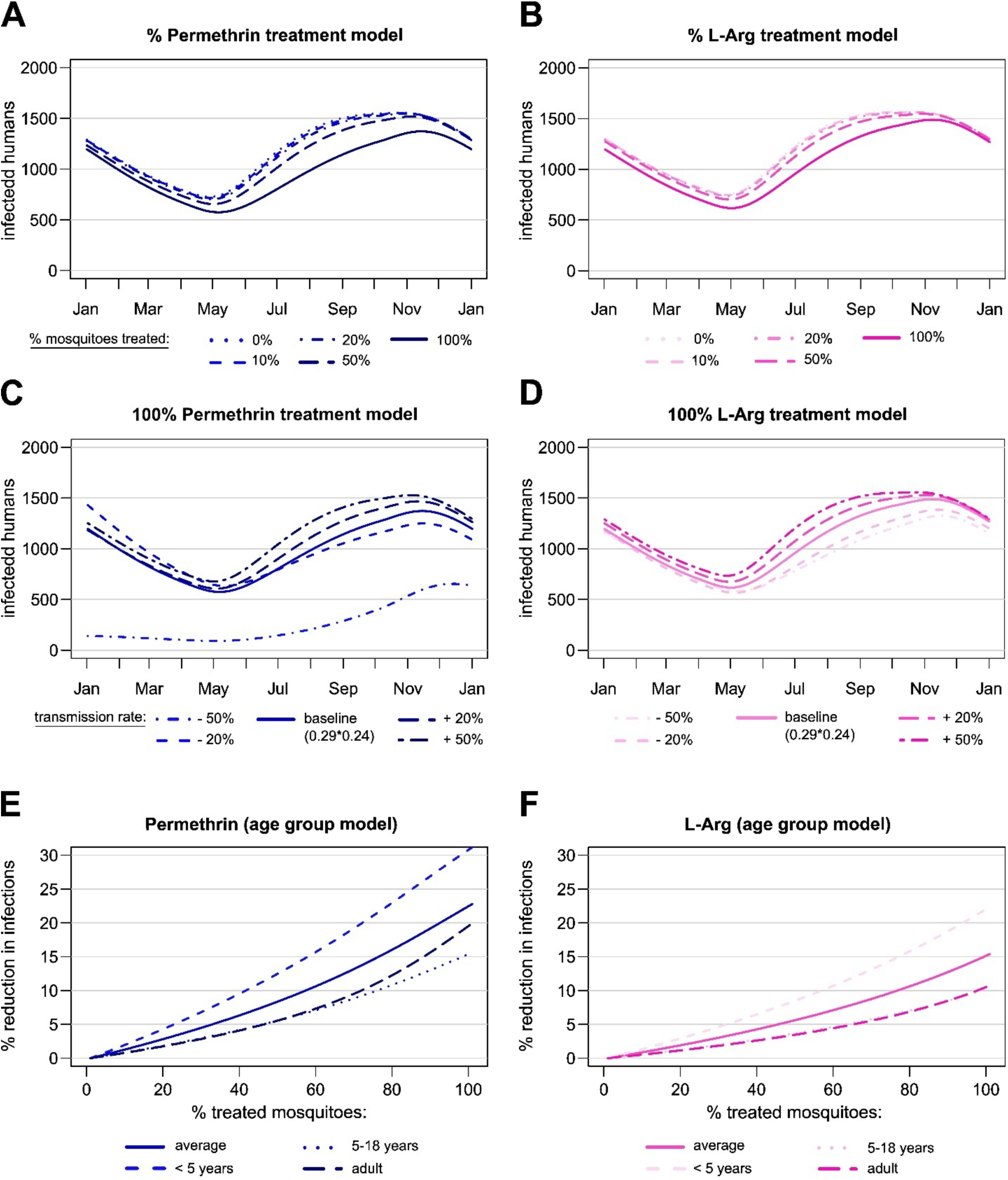
Permethrin and L-Arg reduce human infection rates in compartmental model. (A-B) Human malaria infections (y-axis) when 10%, 20%, 50% or 100% mosquitoes are treated with permethrin (A) or L-Arg (B) compared to the untreated population (baseline model) over the course of one year (y-axis). (C-D) Human malaria infections (y-axis) when 100% mosquitoes are treated with either permethrin (C) or L-Arg (D) over the course of one year (x-axis) assuming different baseline transmission rates (solid line = transmission baseline in all models, dashed and dotted lines = lower (−20%, −50%) and higher (+20%, +50%) baseline transmission). (E-F) Human malaria case reductions (y-axis) per proportion of treated mosquitoes (x-axis) with either Permethrin (E) or L-Arg (F) in different age groups (solid line = average reduction of whole population, dashed and dotted lines = reductions in specific age groups).

Together, these results suggest that sub-lethal permethrin exposure in the absence of additional mortality significantly reduces malaria burden, whilst similar but milder impacts are seen due to RNS augmentation via L-Arg, particularly in young children and low transmission settings.

## Discussion

Owing to their low mammalian toxicity and their fast-acting knockdown effect, pyrethroids remain key to malaria control. All insecticide treated bed nets distributed to date, including the new dual chemistry bed nets, rely on pyrethroids, either alone or in combination with a synergist, insect growth inhibitor/sterilizer or a second insecticide ^80–83^. However, their extensive use has resulted in widespread resistance of mosquito populations across Africa ^84^, raising questions about the continued efficacy of pyrethroids in vector control ^85,86^. Despite their ubiquitous field application, surprisingly little is known about how pyrethroid resistance and/or exposure influences *Plasmodium* development, and whether this has hidden consequences for transmission. Here we demonstrate that pyrethroid exposure of resistant mosquitoes elicits secondary effects, including systemic generation of reactive nitrogen species and a more limited reactive oxygen species response. These shifts in RNS and ROS intrinsically impact *Plasmodium* development, altering the vectorial capacity of both insecticide-resistant and -susceptible mosquitoes. In contrast to prior studies in the mouse model *P. berghei* ^21,24^, our data suggests that ROS may promote parasite development, whilst RNS play a key role in suppressing it. Specifically, we show that both RNS increase through pyrethroid exposure and by oral feeding of L-Arg modulate immune cell composition and trigger production of potent anti-plasmodial effectors to kill invading parasites. Notably, elevated RNS lead to higher levels of ookinete fragmentation and proximal nitration, likely through a non-canonical IMD response thus reducing vectorial capacity. Finally, using an SEIR model we show that permethrin exposure, in the absence of additional mortality, can have a significant impact on parasite transmission, highlighting the importance of their continued use in malaria control.

The upregulation of *NOS* and *CAT* in pyrethroid-resistant mosquitoes, in the absence of a corresponding increase in ROS or RNS likely reflects enhanced redox detoxification mechanisms that maintain redox homeostasis; this aligns with previous studies reporting respiratory changes and oxidative stress in pyrethroid-resistant insects ^5,6,87–89^. Resistance inevitably leads to continued exposure to sub-lethal but substantial doses of insecticides. In mammalian systems, pyrethroid exposure is known to induce a non-specific ROS response ^13^, and in *Anopheles* results in oxidative damage with both type I and II pyrethroids ^90^. In contrast we find that PY-R mosquitoes show minimal ROS perturbation, likely through efficient detoxification of radicals due to the up-regulation of *CAT* and *SOD*. The unexpected systemic and prolonged RNS response has not been previously reported in the context of insecticide exposure. Whilst nitric oxide is a well-established signaling molecule with diverse roles across the animal kingdom ^91,92^, and is essential for development in *Drosophila melanogaster* ^93,94^, it has primarily been studied indirectly in the context of immunity in mosquitoes ^25,95,96^. The observation that pyrethroid exposure leads to a RNS response and consequently a reduced *P. falciparum* infection, combined with the known involvement of nitric oxide in mosquito immunity to *Plasmodium*, suggests that insecticide exposure could influence vectorial capacity, altering malaria transmission dynamics.

Our findings demonstrate that specifically elevated RNS levels in immune cells reduce vectorial capacity by promoting ookinete destruction, likely mediated through granulocyte proliferation and complement activation ^34,97,98^. Whilst RNS manipulation alone leads to changes in haemocyte composition, pyrethroid-treated mosquitoes also exhibited bubbling haemocytes, resembling membrane blebbing during an apoptotic stress response ^99,100^ or possibly the formation of haemocyte-derived microvesicles (HdMVs), which activate TEP1 and thus the mosquito complement system ^34^. Nevertheless, L-Arg-induced immune changes closely resemble those induced by pyrethroid exposure. Haemocyte populations are highly dynamic in response to blood feeding or pathogen challenge in vector mosquitoes ^55,101–104^ and immediate proportional changes had been observed in honeybees following application of the pyrethroid Bifenthrin, similar to what we found after permethrin tarsal exposure ^105^. These data support the idea that direct and long-term shifts in these immune populations drive this lowered infection phenotype. Our RNAseq data further implicate RNS in immune regulation by confirming granulocyte proliferation and upregulation of key immune effectors during ookinete invasion and apoptosis causing caspases, like long caspase (*CASPL2*) ^60,64,73,74^. Granulocytes play a central role in both early and late-stage immunity, particularly against *Plasmodium berghei* ^34^. These phagocytic cells mediate ookinete clearance via the TEP1-dependent complement response through JNK and IMD signaling ^34,39,40,97,106^ and further target early oocysts via JAK-STAT ^97,107,108^. Collectively, our findings describe a previously unrecognized mechanism of how TEP1-dependent lysis of *P. falciparum* can be potentiated by inducing granulocyte nitration. Similar phenomena have only been described in rodent *Plasmodium* models, where hemocyte nitration activated Toll signalling and facilitated TEP1 deposition on oocysts, culminating in the melanisation of mature oocysts ^98^.

Unexpectedly, our RNAseq data revealed downregulation of the JNK, JAK/STAT and TOLL-pathways and including their associated transcription factors *REL1* and *REL2*. In contrast, *IMD*, a key upstream activator of both IMD and JNK signaling, was upregulated. Supporting this, the ubiquitin ligases *DIAP2* and *UEV1A*, which activate IMD via K63-linked ubiquitination, were upregulated, whilst *EFF*, an E2 ubiquitin-conjugating enzyme involved in K48-linked ubiquitination and IMD degradation, was downregulated, together indicating sustained IMD activation ^109,110^. Additionally, the negative IMD regulator *CASPAR* was downregulated ^111^, reinforcing pathway activation. However, multiple downstream factors, including *REL2* were downregulated, whilst key IMD-controlled effectors, such as *TEP1* ^35^, *gambicin* ^70^, *LRIM1* ^69,112^, and *cecropins* ^113^ were overexpressed; this suggests a non-canonical pathway in *Anopheles,* decoupled from upstream JNK and TOLL signaling.

Further supporting an immune-priming effect, fibrinogen and numerous fibrinogen-related proteins (FREPs), such as *ficolin A*, were upregulated following L-Arg treatment. These proteins can hinder *Plasmodium* establishment by forming a dense fibrin network that prevents parasite migration from the blood bolus ^114–116^. Ficolins, in particular, are key pathogen recognition molecules ^117,118^, that may accelerate parasite clearance.

In contrast to RNS levels, continuous elevation of ROS levels in the midgut and haemolymph led to increased oocyst and salivary gland sporozoite numbers, despite no detectable changes in immune cell populations. This may be explained by the downregulation of immune- and midgut-related genes in our RNAseq data, alongside reduced microbial diversity in our 16S analysis that could initiate such a constitutive shut down of immune processes ^119^. Particularly the loss of *Asaia,* which is known to inhibit *Plasmodium,* could be a major contributing factor ^67,120^. Lowered expression of CLIPs is particularly noteworthy, as they could diminish TOLL signaling by preventing Spätzle-mediated activation and blocking melanisation through inhibition of PPOs cleavage to active POs ^121,122^. Furthermore, increased ROS levels could also disrupt epithelial barriers, like septate junctions, in the midgut to increase permeability and facilitate parasite migration as previously seen in multiple mammalian studies where ROS, including H₂O_2_, disrupted epithelial tight junctions to promote infections ^123,124^. In mosquitoes, ROS can also feed back into insulin/IGF-1 signaling and activate phosphorylation of effectors such as FOXO and ERK to lower immunity and promote parasite development ^125^, which could explain the effects seen here. Nevertheless, these effects observed under conditions of systemic ROS elevation do not appear comparable to the transient peaks seen after pyrethroid exposure, and so further studies are needed to determine how ROS in the context of pyrethroid exposure may impact parasite development.

Finally, our modeling simulations indicate that increasing RNS levels through both sub-lethal permethrin exposure or L-arginine feeding could substantially reduce transmission, particularly in low-density vector populations or during periods of reduced mosquito abundance. In particular, young children and infants, who are most vulnerable to infection, can benefit the most. Similar mathematical modelling approaches for other transmission settings across Africa are scarce, but one hypothesized that 80% IRS coverage with bendiocarb could almost completely interrupt transmission ^126^ and ITN usage of 80% in a low-transmission setting could reduce malaria transmission below 1% ^127^, closely mirroring an intervention based around permethrin. Although these models closely mimic the interruption in transmission, it is difficult to directly compare them due to differences between SEIR and models utilising entomological inoculation rates ^128,129^ and those that consider only known insecticidal impacts. Interestingly, when considering the transmission blocking impact of permethrin, models showing that low-level insecticide resistance would likely increase malaria incidence due to failure of insecticide-based tools could be incorrect^130^. Whilst our simulations cannot fully capture real-world settings, such as the precise uptake levels, concurrent interventions or geographic heterogeneity, the permethrin-based model demonstrates a clear transmission blocking impact of pyrethroid insecticides beyond mosquito lethality which should be considered in future modelling strategies.

Our study reveals a critical intersection between pyrethroid resistance and mosquito immunity, demonstrating that reactive nitrogen species, induced by pyrethroid exposure, lead to proliferation of granulocytes and activation of effector molecules against *Plasmodium falciparum*. These findings position redox homeostasis as a previously underappreciated but potentially exploitable target for vector control. Importantly, we provide the first molecular evidence that pyrethroids modulate mosquito immunity, influencing parasite transmission. Together, our results highlight the broader biological impacts of pyrethroid exposure, beyond direct lethality, and underscore the need to fully consider these effects when evaluating the future role of pyrethroids in malaria control strategies.

## Resource availability

All experimental data produced in this study are outlined in the text, figures, supplementary information and tables and supplementary figures. The sequenced transcriptomic data has been deposited in SRA under accession PRJNA1212997.

## Acknowledgements

This study was funded through the Deutsches Zentrum für Infektionsforschung (DZIF, TTU03.705 (VAI) and TTU03.708 (RK)) and an ERC Starting Grant (101075634, ReMVeC) awarded to VAI and Deutsche Forschungsgemeinschaft (DFG, German Research Foundation) – project number 240245660-SFB 1129, MD program to VAI for the support of LH’s medical thesis. We thank Liverpool Insect Testing Establishment (part of iiDiagnostics Ltd.) for providing Kisumu and The Centre National de Recherche et de Formation sur le Paludisme for providing Tiefora to VAI. We thank the Infectious Diseases Imaging Platform (IDIP) in Heidelberg for advice on microscopy experiments and provision of the SP8 microscope. We also thank Stephanie Blandin, French Institute of Health and Medical Research for providing the TEP1 antibody and Teun Bousema, Radboud University Nijmegen for the Pfs25 antibody and providing NF54. Finally, we thank Hilary Ranson, Freddy Frischknecht and Franziska Hentzschel for helpful comments on the manuscript.

## Author contributions

PH was involved in all experiments carried out, either directly performing the experiments or supervising students working on this project. CW assisted with imaging optimization, explored haemocyte bubbling and assisted with nitrotyrosine imaging. LH performed H_2_O_2_ and dsCAT experiments, including the infectious feeds and 16S generation, in addition to assisting with LC_50_ generation. DS and AB performed ROS haemocyte imaging and optimized RNAi of the ROS-related enzymes, and dot-blot and egg counting respectively. LD worked under the supervision of RK to produce the modelling data. JBM reared the mosquitoes. VAI conceived of, funded and supervised the project throughout and analysed the RNAseq data. PH and VAI drafted the manuscript.

## Declaration of Interests

The authors declare no competing interests in this paper.

## Supplemental figures

**Figure S1.**
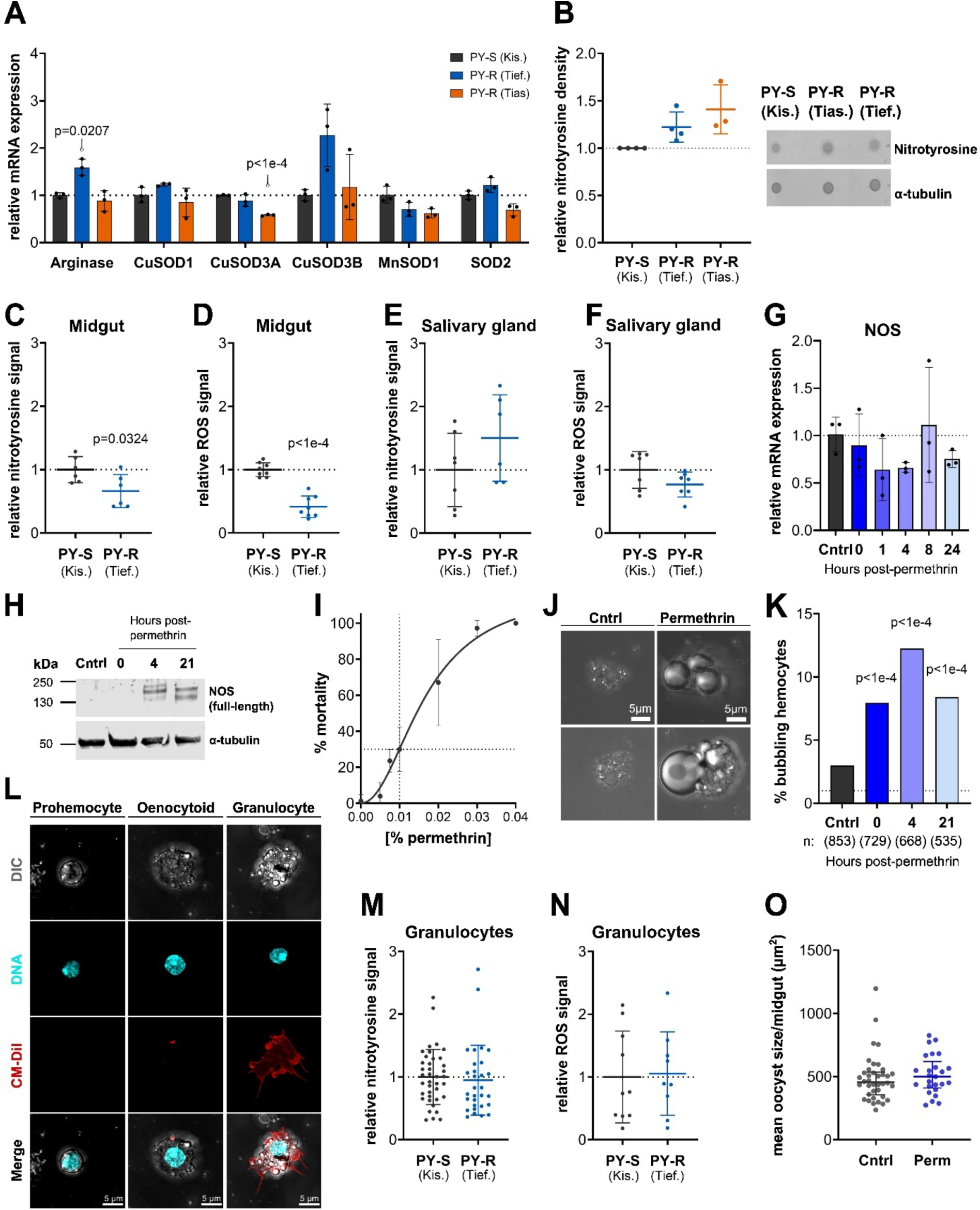
ROS and RNS are decreased in PY-R midguts while permethrin affects NOS and haemocytes, related to Figures 1 and 2. (A) Relative mRNA expression levels (y-axis) of arginase and superoxide dismutases (SODs) in two pyrethroid-resistant (PY-R) populations (Tiassalé – Tias., Tiefora – Tief.) compared to a susceptible (PY-S) population (Kisumu – Kis.) (x-axis). Significance: t-test with Bonferroni correction. (B) Representative dot blot (right) showing nitrated proteins (nitrotyrosines) and *α*-tubulin (internal loading control) in both PY-R compared to PY-S mosquitoes. Quantitative analysis of the dot blot (left) showing relative density normalized to the internal loading control (y-axis) in both PY-R compared to PY-S mosquitoes (x-axis). Significance: not significant with a t-test using paired ratios. (C-F) Quantitative analysis of the nitrotyrosine (C and E) and ROS (D and F) fluorescence signal (y-axis) in dissected midguts and salivary glands of PY-R (Tiefora) relative to PY-S (x-axis). Error bars show standard deviation; each point is one midgut or salivary gland. Significance: t-tests. (G) Relative mRNA expression (y-axis) of NOS at multiple timepoints (0-to 24 hours) following 0.75% permethrin exposure (1-hour) in PY-R Tiefora (x-axis). Significance: one way ANOVA with Dunnet’s test. (H) Western blot showing NOS protein and *α*-tubulin (internal loading control) at three timepoints (0-, 4- and 21 hours) after 0.01% permethrin tarsal contact of pools of five PY-R mosquitoes for 30 min (x-axis). (I) Insecticide-induced mortality (y-axis) to increasing permethrin concentrations in acetone (x-axis) of PY-R. Error bars show standard deviation. Data pooled from 6-7 individual biological replicates of at least ten mosquitoes per concentration in each rep. Intersection of dotted lines shows LC_30_ concentration (0.01%). (J) Confocal brightfield images of extracted granulocytes after 30 min tarsal exposure to 0.01% permethrin in acetone or to acetone-only (Cntrl). Scale bar = 5 µm. (K) Percentage of, bubbling’ haemocytes out of total number of counted haemocytes (0-, 4- and 21 hours) after tarsal contact to 0.01% permethrin (x-axis). n = counted cells per timepoint. Significance: chi-square test. (L) Confocal images classifying prohaemocyte, oenocytoid and granulocyte. Gray = DIC/brightfield, cyan = DNA staining (Hoechst), red = cell marker (Vybrant CM-Dil). Scale bar = 5 µm. (M and N) Quantitative analysis of the nitrotyrosine (M) and ROS (N) fluorescence signal (y-axis) in extracted granulocytes of PY-R (Tiefora) relative to PY-S (x-axis). Error bars show standard deviation; each point is one granulocyte. Significance: not significant with a t-test (log transformations for normality done in M). (O) Mean oocyst size per midgut in PY-R at 5 days pIBM in permethrin 0.01% treated (Perm) or acetone-only treated (Cntrl) (y-axis). Each point is average oocyst size in one midgut. Significance: not significant with a t-test (log transformation for normality performed).

**Figure S2.**
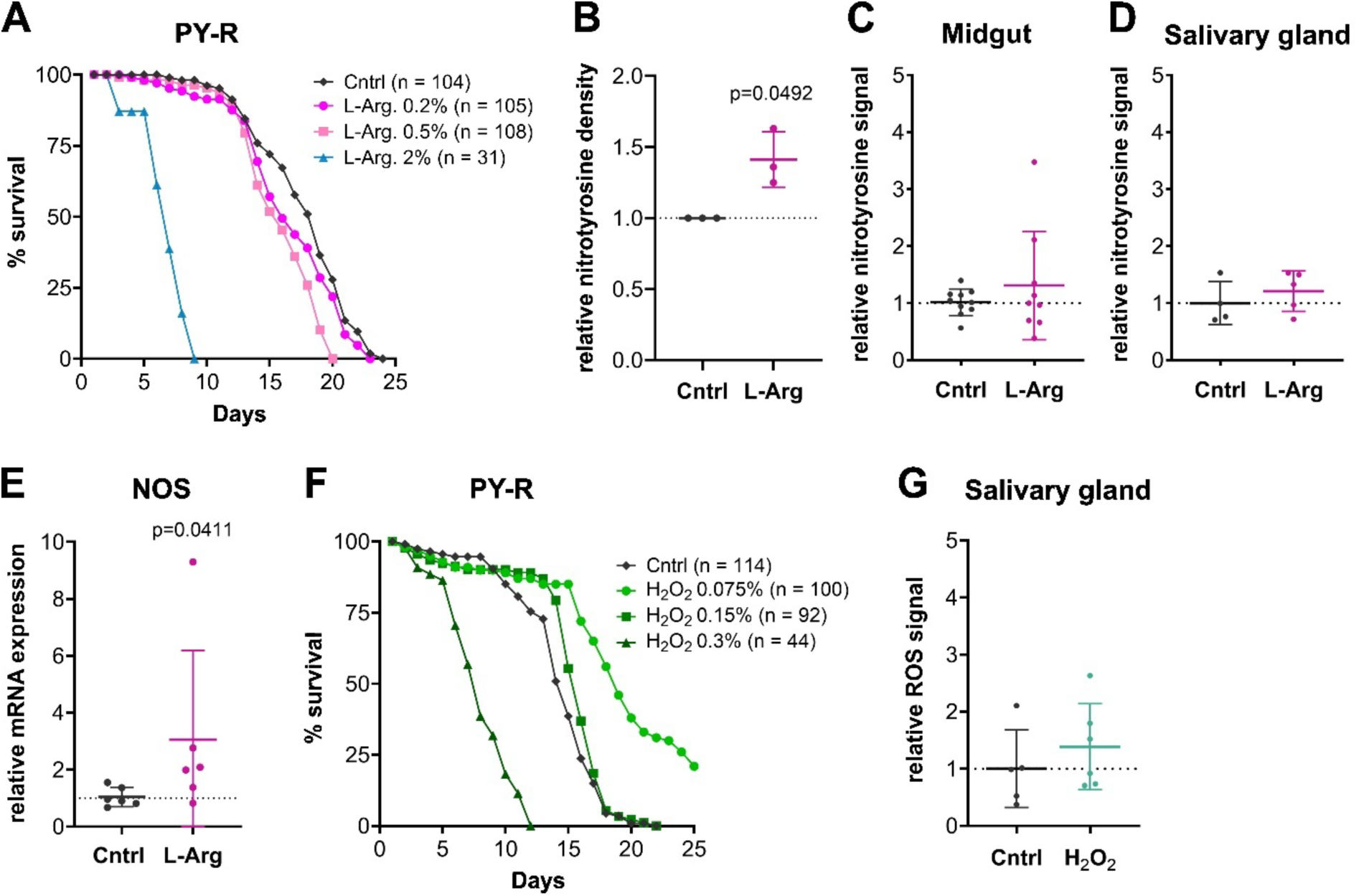
L-arginine specifically affects NOS/RNS in immune cells whilst hydrogen peroxide increases ROS systemically, related to Figure 3. (A) Mosquito survival (y-axis) per day (x-axis) in PY-R receiving Cntrl or L-Arg supplemented diet. n = number of mosquitoes per tested concentration or control. (B) Quantitative dot blot analysis showing relative nitrotyrosine density normalized to the internal control (y-axis) in PY-R receiving Cntrl or L-Arg treatment (x-axis). Significance: t-test with paired ratios. (C and D) Quantitative analysis of the nitrotyrosine fluorescence signal (y-axis) in dissected midguts (C) and salivary glands (D) of L-Arg treated PY-R (Tiefora) relative to Cntrl treatment (x-axis). Error bars show standard deviation; each point is one midgut or salivary gland. Significance: not significant with a t-test. (E) Relative mRNA expression (y-axis) of NOS in PY-R Tiefora receiving L-Arg or Cntrl treatment (x-axis). Significance: Mann Whitney test. (F) Mosquito survival (y-axis) per day (x-axis) in PY-R receiving Cntrl or H_2_O_2_ supplemented diet. n = number of mosquitoes per tested concentration or control. (G) Quantitative analysis of the ROS fluorescence signal (y-axis) in dissected salivary glands of H_2_O_2_ treated PY-R (Tiefora) relative to Cntrl treatment (x-axis). Error bars show standard deviation; each point is one salivary gland. Significance: not significant with a t-test.

**Figure S3.**
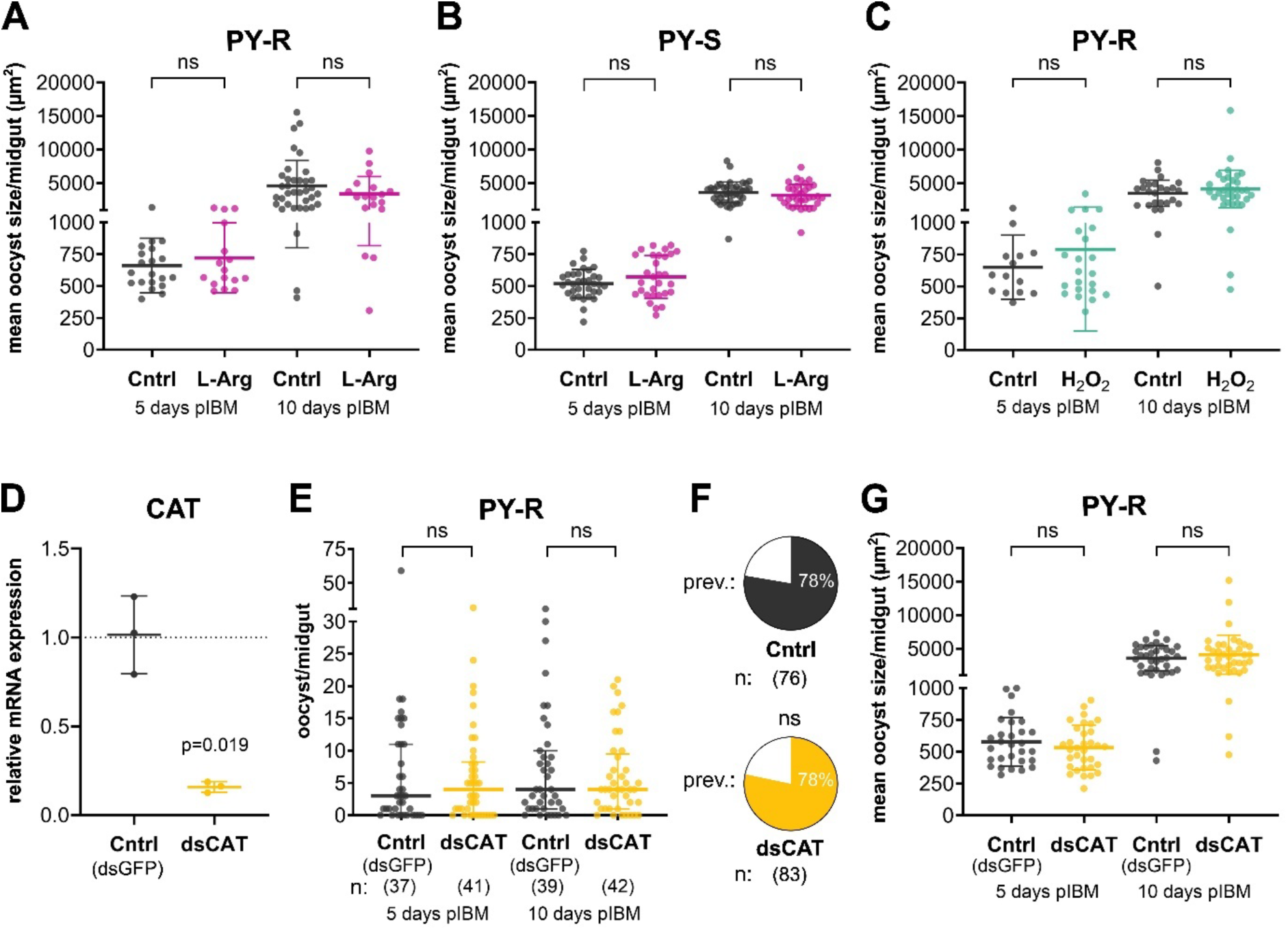
RNS and ROS increase does not affect oocyst maturation, related to Figure 4. (A - C) Mean oocyst size per midgut in PY-R and PY-S (y-axis) at 5 days pIBM and 10 days pIBM receiving either Cntrl, L-Arg (A and B) or H_2_O_2_ (C) treatment (x-axis). Each point is average oocyst size in one midgut. Significance in A and B: all non-significant through t-tests (log transformations for normality performed). Significance in C: all non-significant through Mann Whitney test. (D) Relative mRNA expression levels of CAT (y-axis) in PY-R following injection of dsRNA against CAT (dsCAT) compared to GFP as control (dsGFP) (x-axis). Error bars show standard deviation; each point is one biological replicate of seven pooled mosquitoes. Significance: t-test. (E) Oocysts per midgut in PY-R at 5 days pIBM and 10 days pIBM (y-axis) injected 3 days prior to infection with dsCAT or dsGFP (Cntrl) (x-axis). Each point is oocyst count of one midgut of three individual infectious feeds. Significance: all non-significant with t-tests (log transformations for normality performed). (F) Infection prevalence combining data from both 5 days pIBM and 10 days pIBM. N = total number of midguts per treatment. Significance: non-significant with a chi-square test. (G) Mean oocyst size per midgut in PY-R at 5 days pIBM and 10 days pIBM (y-axis) injected 3 days prior to infection with dsCAT or dsGFP (Cntrl) (y-axis). Each point is average oocyst size in one midgut. Significance: non-significant with a t-test (log transformations for normality performed).

**Figure S4.**
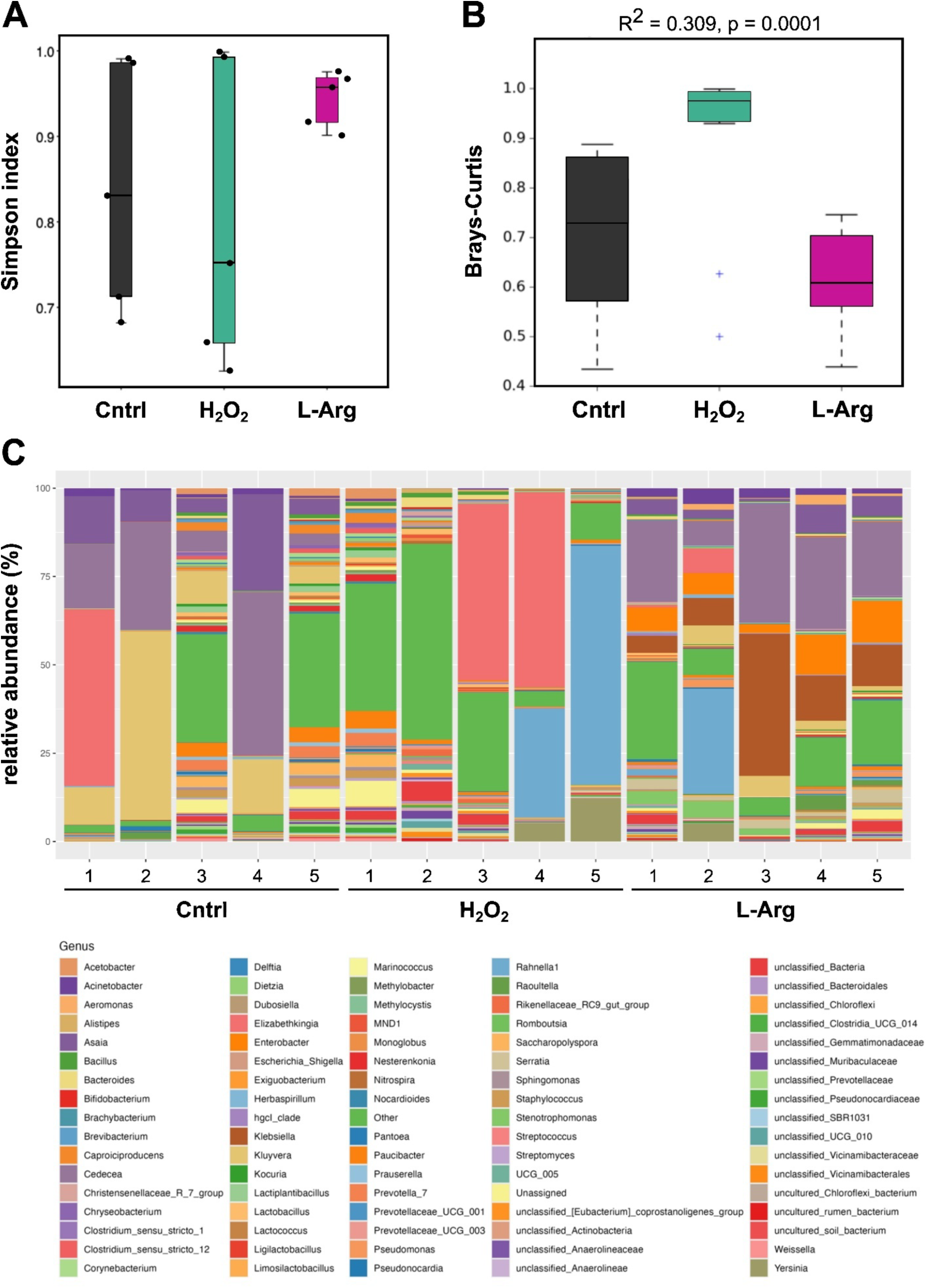
Hydrogen peroxide but not L-arginine decreases bacterial diversity and eradicates Plasmodium-inhibiting species, related to Figure 4. (A) Simpson Index showing alpha diversity (y-axis) in Cntrl, H_2_O_2_ and L-Arg treated PY-R mosquitoes (x-axis). Each dot represents pools of five whole mosquitoes. (B) PERMANOVA analysis in a Brays-Curtis plot showing beta diversity for calculated abundances (y-axis) in Cntrl, H_2_O_2_ and L-Arg treated PY-R mosquitoes (x-axis). (C) Relative abundance plot in % (y-axis) for each pool of five PY-R individuals in Cntr, H_2_O_2_ or L-Arg treatment groups (x-axis) at genus level.

**Figure S5.**
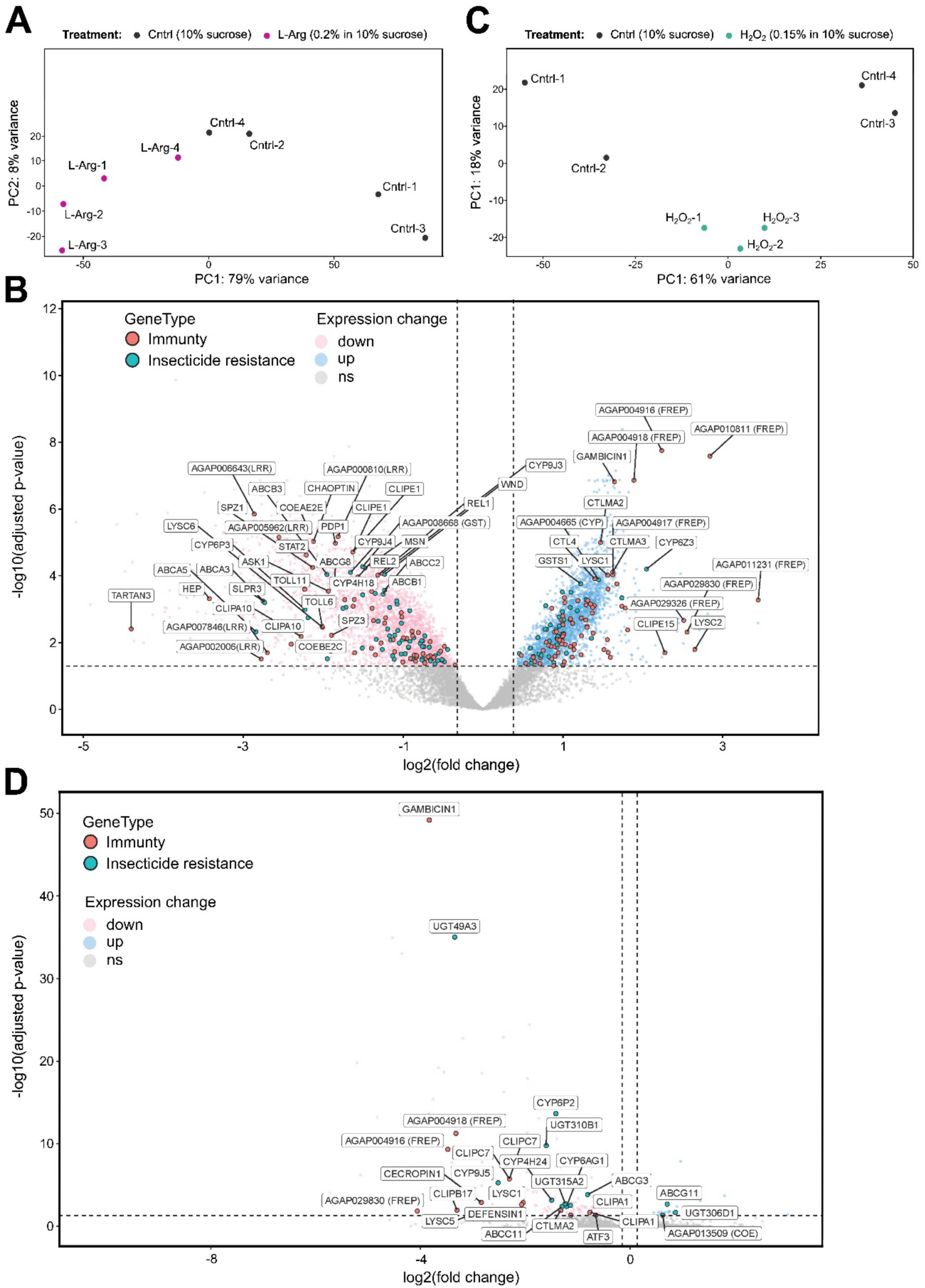
L-arginine and hydrogen peroxide lead to differential expression of immune- and resistance related genes, related to Figure 5. (A) Principal component analysis (PCA) showing PC1 with 79% variance (x-axis) and PC2 with 8% variance between Cntrl or L-Arg treated PY-R mosquitoes. Each point is one biological replicate of 6 mosquitoes. (B) Volcano plot showing significantly up- and down-regulated immunity-(red) and insecticide resistance-related (blue) genes in L-Arg treatment 21 hours post infection with log2 fold change (x-axis) and adjusted p-value (y-axis). (C) Principal component analysis (PCA) showing PC1 with 61% variance (x-axis) and PC2 with 18% variance between Cntrl or H_2_O_2_ treated PY-R mosquitoes. Each point is one biological replicate of 6 mosquitoes. (D) Volcano plot showing significantly up- and down-regulated immunity-(red) and insecticide resistance-related (blue) genes in H_2_O_2_ treatment 21 hours post infection with log2 fold change (x-axis) and adjusted p-value (y-axis).

**Figure S6.**
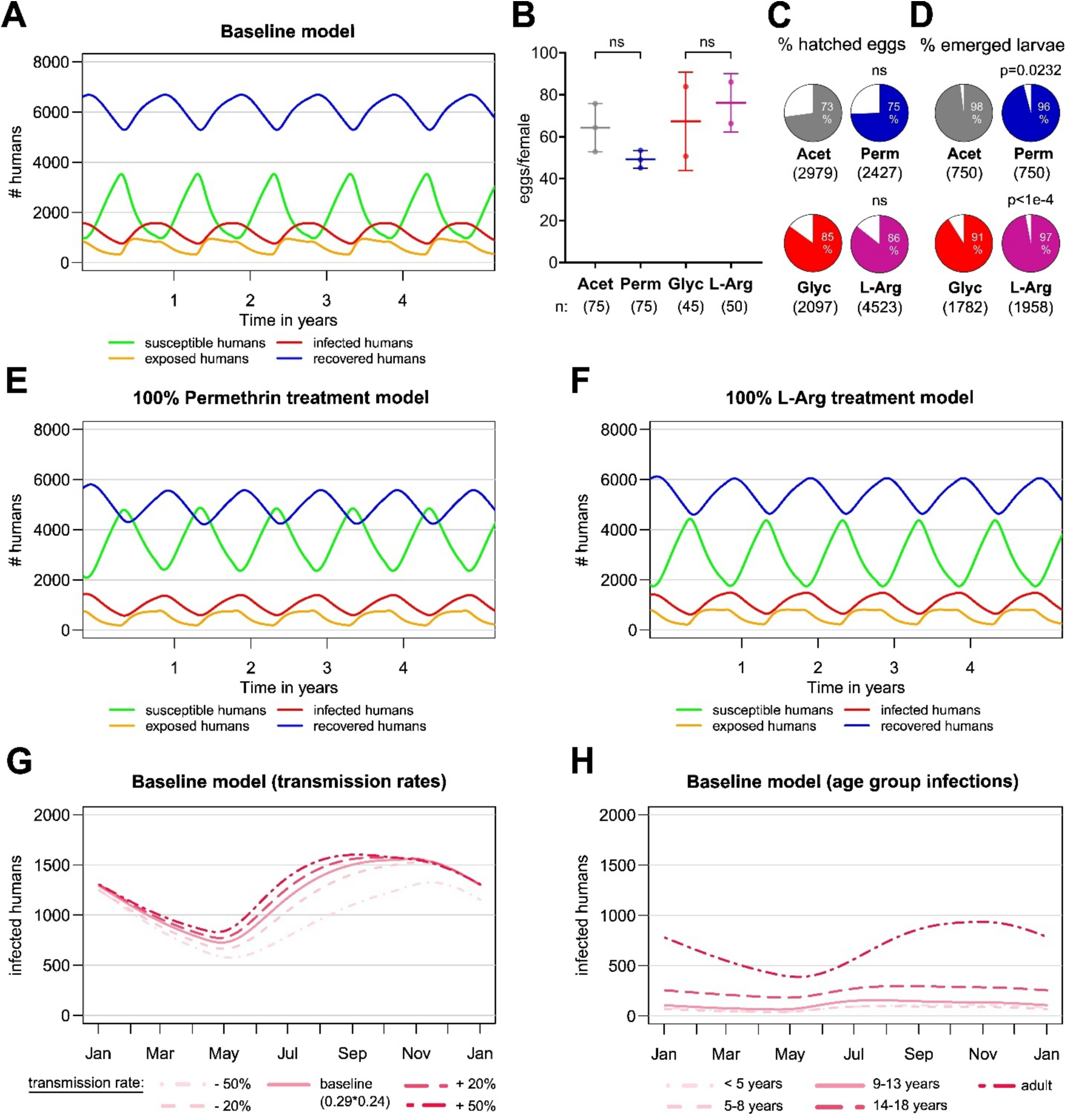
SEIR model shows permethrin and L-Arg decrease the infected human population, related to Figure 6. (A) Naïve SEIR model showing susceptible-, exposed, infected and recovered humans (y-axis) over the course of multiple years in a setting in southern Burkina Faso (x-axis). (B) Average number of eggs produced per PY-R female (y-axis) in Perm, Acet, Glyc or L-Arg treated PY-R mosquitoes (x-axis). n = total number of females per treatment. Error bars show standard deviation; each point is average egg number/mosquito of at least 20 pooled females. Significance: non-significant with a t-test. (C-D) Percentage of hatched mosquito eggs (C) and emerged larvae (D) when PY-R adult generation was treated with Perm, Acet, Glyc or L-Arg n = number of total investigated eggs or larvae per treatment. Significance: chi-square test. (E-F) Adaptation of naïve SEIR model for 100% treatment of mosquitoes with permethrin (E) or L-Arg (F) showing susceptible-, exposed, infected and recovered humans (y-axis) over the course of multiple years in a setting in southern Burkina Faso (x-axis). (G) Human malaria infection in the naïve model (y-axis) over one-year (x-axis) assuming different baseline infection rates (solid line = base used for following models, dashed and dotted lines = lower (−20%, −50%) and higher (+20%, +50%) probabilities of infectious bites). (H) Human malaria infections (y-axis) over one year (x-axis) per age group in naïve model.

## STAR methods

### Contact for Reagent and Resource sharing

Further information and requests for resources and reagents should be directed to and will be fulfilled by the Lead Contact, Victoria Anne Ingham when possible.

### Experimental models

#### Rearing of Anopheles gambiae s.s and Anopheles coluzzii mosquitoes

All mosquitoes were reared under standard insectary conditions at 27°C and 70-80% humidity in a 12-hour light-dark cycle with 1 hour dawn:dusk. Adults were provided 10% sucrose solution in *ad libitum* and where treated, supplemented with 0.2% L-arginine (L-Arg), 0.2% Glycine (Glyc) or 0.15% hydrogen peroxide (H_2_O_2_). Mosquitoes received reconstituted human blood (Blood bank, University hospital Heidelberg Germany) for egg production to sustain the following generations using a Hemotek® membrane feeding system. Larvae received ground fish food (Tetramin, Germany) twice per day. Insecticide resistant mosquito populations were selected on 0.05% deltamethrin and 0.75% permethrin every fourth generation. Pyrethroid resistant mosquitoes (PY-R) used were Tiefora (*An. coluzzii*) originally colonized from the Cascades region in Burkina Faso by Liverpool School of Tropical Medicine (LSTM) and Centre National de Recherche et de Formation sur le Paludisme ^131^ and Tiassalé (*An. gambiae*) originally colonized from Côte D’Ivoire by Centre Suisse de Recherches Scientifics en Côte D’Ivoire ^132^. The insecticide susceptible (PY-S) Kisumu strain (*An. gambiae*) was used as the comparator ^133,134^. All experiments were performed with adult mosquitoes aged between 2-6 days and all mosquitoes have been reared at Heidelberg University, Medical Faculty.

#### Culturing of Plasmodium falciparum NF54 parasites

Asexual parasites of the NF54 strain of *P. falciparum* (Nijmegen, Netherlands) were maintained between 0.5% and 3% parasitemia at 37°C in human erythrocytes (O^+^) with 2.5% total hematocrit (Blood bank, University hospital Heidelberg, Germany). Medium used was RPMI 1640 (Corning, Manassas, VA) supplemented with 25mM HEPES, 10mg/l hypoxanthine, 0.3% sodium bicarbonate and 10% filtered and heat-inactivated human serum (Haema AG, Leipzig, Germany). The gas mixture was 5% O_2_, 5% CO_2_ and balanced N_2_ according to established protocols ^135,136^. Gametocyte cultures were induced by splitting asexual cultures to 0.3% in 5% hematocrit and incubated for 16-18 days with daily media change without disturbance of the blood layer to produce mature stage V male and female gametocytes.

## Method details

### RNA extraction, cDNA synthesis and qPCR analysis

RNA from whole bodies of pools of 5-7 individual female mosquitoes of various treatment and control groups was extracted with the Arcturus PicoPure RNA isolation Kit (Thermo Fisher Scientific, Germany) in triplicates by homogenization in 100 μl extraction buffer and following of the manufacturer’s instructions. DNA contamination was removed using the RNase-free DNase set (Qiagen GmbH, Germany) following previously published protocols ^137^. First-strand cDNA synthesis was performed with the SuperScript® III Reverse Transcriptase protocol (Thermo Fisher Scientific, Germany) using Oligo(dT)20 primers to select for messenger RNA. For subsequent purification of the cDNA product the Qiagen QIAquick® DNA purification Kit was used according to the manufacturer’s instructions (Qiagen GmbH, Germany) and the resulting dsDNA was quantified using a NanoDrop Lite spectrophotometer (Thermo Fisher Scientific, Germany). Primers with lengths between 15-20bp were designed with Primer Blast (NCBI) ^138^ (Table S1) for quantitative real-time PCR (qPCR). Exon-exon junctions were spanned where possible for products of 80-150bps with 40-60% GC content and potential off-target binding checked in VectorBase. Brilliant III Ultra-Fast SYBR® Green qPCR Master Mix was used (Agilent, Germany) on a CFX96™ Real-Time System (Bio-Rad, Germany) that includes the CFX Maestro 1.1 software (Bio-Rad, Germany). All cDNA samples were diluted to a concentration of either 2 ng/μl (*SODs*), 4 ng/μl (*CAT*, *Arginase*) or 8 ng/μl (NOS) depending on the expression levels of the respective target gene. Reactions were prepared in triplicates by adding 10 μl 2x SYBR master mix, 0.3 μM of primers and molecular grade H_2_O to the diluted cDNA to make 20 μl reactions. The housekeeping genes: *elongation factor Tu (EF)* (Gene ID: *AGAP005128*) and 40S *ribosomal protein S7 (S7)* (Gene ID: *AGAP010592*) were used for normalization of the relative gene expression (Table S1). The temperature protocol for qPCRs was 3 minutes at 95°C for initial denaturation, followed by 40 cycles of 10 seconds at 95°C and 10 seconds at 60°C. This method was used to determine expression of *NOS (AGAP029502)*, *CAT (AGAP004904*), mitochondrial superoxide dismutases *MnSOD1 (AGAP010517)* and *CuSOD2 (AGAP005234)*, cytoplasmic *CuSOD1 (AGAP007497)*, extracellular *CuSOD3A (AGAP010347-RA)* and *CuSOD3B (AGAP010347-RB)* and *Arginase* (*AGAP008783*) (Table S1).

### Immuno-dot blotting

Protein extracts of whole mosquitoes (see section Western Blotting) were denatured for 10 min at 70°C with 4x Laemmli buffer and 1 M DTT and exactly 1 μl per sample was pipetted onto 0.2 μm nitrocellulose membranes (Bio-Rad), blocked for 1 hr in milk-PBS-T blocking buffer, washed 3 times for 20 min in PBS-T and incubated for 1 hr in polyclonal rabbit anti-nitrotyrosine primary antibody (1:1000) in blocking buffer (Thermo Fisher Scientific, Germany). This was followed by three 20 min washing steps in PBS-T and incubation for 2 hr in secondary donkey anti-rabbit 800 (1:15000) and donkey anti-mouse 700 (1:20000) antibodies (Li-Cor) supplemented with 0.01% SDS at RT. After washing membranes three more times in PBS-T and 1x in PBS images were acquired with LI-COR Odyssey CLx imaging system. Images were processed and intensity of dots analysed quantitatively with the Image Studio software by normalizing the signal of nitrotyrosine to the internal control (α-tubuline).

### Confocal microscopy methods and quantitative analysis

Point laser scanning confocal microscopy was performed on a Leica SP8 microscope TCS DLS Confocal and SPIM (Leica Microsystems, Wetzlar, Germany) using HC PL APO CS2 63x/1.4 N.A. oil immersion objective equipped with PMT and HyD detectors. Images were acquired sequentially and with variable z-stacks with, depending on the tissue thickness, 30 μm to 60 μm z-stack size. Laser excitation was UV 405 nm (DNA), Argon 488 nm (Pfs25), DPSS Yellow 561 nm (Nitrotyrosine) and HeNe 633 nm (Nitrotyrosine, ROS). Brightfield images were obtained from a transmitted light PMT detector.

For mosquito midgut ROS and nitrotyrosine imaging three individual fields of view on the tissue were randomly selected with the UV 405 nm light and imaged. Per salivary gland 1-3 spots in total from the middle lobe and from distal lateral lobes were randomly selected with UV 405 nm laser and imaged. Quantitative analysis was performed in Fiji by comparing fluorescent signal of either nitrotyrosine or ROS against the respective control per experiment. For midguts, integrated density was measured for each stack through one midgut cell layer of all three selected spots and averaged. The change in integrated density of all treatment groups was compared to the average of the respective controls per experiment. The method of corrected total cell fluorescence (CTCF) in Fiji was used for quantification of salivary gland fluorescent signal through all stacks of one salivary gland cell layer and through whole individual haemocytes and compared to the average of the respective control per experiment ^57^.

### Immunofluorescence assay of midguts, salivary glands

Midguts were prefixed in 4% PFA in PBS for 1 min and opened longitudinally on ice-cold PBS. After fixation for 45 min in 2% PFA at RT, midguts were washed with PBS and blocked for at least 1 hr in a buffer containing 1% BSA, 0.1% Triton X-100 and PBS (PBST). The midguts were incubated overnight with polyclonal rabbit anti-nitrotyrosine primary antibody (Thermo Fisher Scientific, Germany) 1:1000 in PBT at 4°C, followed by washing and incubation with either Alexa Fluor™ 568 or Alexa Fluor™ 647 goat anti-Rabbit IgG secondary antibody (Thermo Fisher Scientific, Germany) for 3-4 hr at RT. Nuclei were counterstained with 20 μM Hoechst 33342 (Thermo Fisher Scientific, Germany) for 30 min at RT, followed by washing and mounted with Vectashield® mounting medium (VWR, Cat. No.: VECTH-1000) on glass slides for confocal microscopy analysis. Mosquito salivary glands were seeded on glass bottom dishes (MaTek Corporation, Ashland, USA) coated with poly-D-lysine (Gibco, Germany), fixed for 1hr in 4% PFA, blocked for at least 1 hr in PBST at RT and incubated overnight at 4°C with rabbit anti-nitrotyrosine primary antibody (Thermo Fisher Scientific, Germany). All further steps were similar to treatment of midguts, excluding the mounting step as salivary glands were imaged on the bottom of the glass dish in PBS.

### ROS staining of midguts, salivary glands

Dissected tissues were incubated in a solution containing 5μM CellROX™ Deep Red (Invitrogen, Germany) and 20μM Hoechst for 30 min at RT, washed with PBS and fixed for 1hr in 2% PFA (midguts) or 4% PFA (salivary glands). After washing, midguts and salivary glands were mounted on glass slides with Vectashield® mounting medium (VWR) and tissues were imaged not longer than 2 hours after fixation.

### Immunofluorescence assay of haemocytes

IFAs of haemocytes (granulocytes) were performed in μ-Slide 8 well glass bottom dishes (ibidi, Fitchburg, USA) previously coated for 20 min at 28°C with 5 mg/ml concanavalin A in H_2_O. Concanavalin A was removed and wells rinsed with PBS before the staining was performed. Haemolymph of pools of 25-30 mosquitoes was extracted by puncturing of mosquito thorax and 8 min centrifugation at 3000 g. Cells were resuspended in Schneider’s *Drosophila*-Medium (Thermo Fisher Scientific, Germany) and seeded on the dish for 30 min at 28°C in the dark. Wells rinsed with PBS twice and cells were fixed with 4% PFA for 20 minutes at 28 °C. After washing twice and blocking for 1 hr in PBT antibody staining was performed by adding polyclonal rabbit anti-nitrotyrosine primary antibody (Thermo Fisher Scientific, Germany) 1:1000 for 1 hr at RT. Cells were washed three times in PBT and incubated with Alexa Fluor™ 647 goat anti-Rabbit IgG secondary antibody and 20μM Hoechst and again washed three times. Imaging was performed on the glass bottom dish in PBS.

### ROS staining of haemocytes

ROS staining of haemocytes was performed in μ-Slide 8 well glass bottom dishes (ibidi, Fitchburg, USA) previously coated for 20 min at 28°C with 5 mg/ml concanavalin A in H_2_O. Concanavalin A was removed and wells rinsed with PBS before the staining was performed. Extracted haemolymph was incubated in CellROX™ Deep Red staining (Invitrogen, Germany) in a working concentration of 5 μM and 20 μM Hoechst for 30 min at 28°C in the dark during the seeding process on the glass bottom dishes. Supernatant was carefully discarded, wells rinsed with PBS twice and cells were fixed with 4% PFA for 20 minutes at 28 °C. After washing twice, cells were imaged on the glass bottom dish in PBS.

### Western blotting

Whole protein was extracted from pools of 5 adult females or haemolymph of 25-30 females in freshly prepared protein extraction buffer (PEB) containing 0.1% sodium dodecyl sulphate, 150 mM sodium chloride, 1% Triton X-100, 5 mM EDTA, 20 mM Tris-HCl (pH 7.4) and 1X protease inhibitor cocktail (Roche, Basel, Switzerland). Haemolymph was extracted by puncturing of mosquitoe’s thorax and 8 min centrifugation at 3000 g. Samples were homogenized, briefly sonicated at 60% for ten intervals and left on ice for 30 min before removing the insoluble parts by centrifugation for 10 min at 4°C. Protein concentration was quantified by Bradford assay as previously described by using Pierce™ Coomassie Protein Assay Reagent ^139^ (Thermo Fisher Scientific, Germany) and a standard plate reader. The extracts were diluted in PEB to 25 ng, mixed with 4x Laemmli buffer and 1 M DTT, denatured at 70°C for 10 min, loaded onto 4-20% Mini-PROTEAN® TGX Stain-Free™ protein gels (Bio-Rad, Germany) and run at 100 V for 40-50 min in a Mini-PROTEAN Tetra Vertical Electrophoresis Cell with NuPAGE MES SDS Running Buffer (Thermo Fisher Scientific, Germany). Proteins were transferred onto 0.2 μm nitrocellulose membranes (Bio-Rad, Germany) using Trans-Blot Turbo Transfer System. Successful transfer was confirmed using Ponceau S staining (Thermo Fisher Scientific, Germany). Blocking was performed in milk-PBS-T blocking buffer (5% milk powder, 0.1% Tween® 20 for 1 hr at RT while shaking. Membrane was incubated on a shaker with primary antibodies in milk-PBS-T blocking buffer, including affinity purified rabbit anti-uNOS Polyclonal Antibody (Thermo fisher scientific, Germany) (1:500), Rabbit polyclonal anti-catalase Antibody (Proteintech, Rosemont, USA), Rabbit polyclonal anti-Tep1 Antibody ^35^ and internal control mouse anti-α-Tubulin monoclonal antibody (Sigma Aldrich, Germany) (1:2000) overnight at 4°C. After three washing steps in PBS-T membranes were incubated in fluorophore-conjugated secondary donkey anti-rabbit 800 (1:15000) and donkey anti-mouse 700 (1:20000) antibodies (Li-Cor) supplemented with 0.01% SDS for 1hr in the dark at RT while gently shaking. Membranes were then washed 3x 5 min in PBS-T, 15 min in PBS-T and 5 min in PBS at RT on a shaker before image acquisition at the LI-COR Odyssey CLx imaging system. Images were processed and bands analysed quantitatively with the Image Studio software by normalizing the signal to the internal control (α-tubulin).

#### Dose-response assay to permethrin for LC_30_ determination

Dose-response assays to increasing doses of permethrin were performed to establish an LC_30_ dose to permethrin. Permethrin (PESTANAL®, Merck, Germany) was diluted in acetone to concentrations ranging from 0.005% to 0.04% for the assay with PY-R Tiefora mosquitoes. 0.5 ml of diluted insecticide was applied to the plate and 0.5 ml acetone added for proper dispersion. Plates were left on a shaker at RT for at least 4 hr until acetone was completely evaporated and stored in the fridge for up to two weeks or until three times used. For the exposures mosquitoes were aspirated onto plates through a small hole in the plastic lid and sealed with parafilm. Exposure time was 30 min in an incubator with standard insectary conditions before mosquitoes were released in a cage, aspirated in individual cups and mortality scored 24 hr later.

### Visual haemocyte classification

Haemocytes were visually classified by confocal imaging into the three major classes, prohaemocytes, oenocytoids and granulocytes, via size and morphology by confocal microscopy as previously described ^54,106^. The haemocyte specific dye Vibrant CM-Dil was used to additionally confirm classification as granulocytes initially. All counting experiments were performed in triplicates and for each replicate at least 50 haemocytes were counted per treatment.

### P. falciparum infections of An. gambiae s.s and An. coluzzii mosquitoes

Female mosquitoes were fed on prewarmed glass membrane feeders with NF54 gametocyte culture and diluted with reconstituted blood to 0.2% stage V gametocytemia. For permethrin exposure experiments females were put on glass petri dishes coated with 0.01% permethrin in acetone or acetone-only for 7.5 min 3 hours before the infectious feed. In all experiments unfed and not fully engorged mosquitoes were removed in a custom-made glove box approximately 24-26 hours post infectious blood meal into 15ml falcon tubes filled with 70% ethanol and frozen in −80°C. Remaining mosquitoes received 10% sucrose *ad libitum*, L-Arg- or H_2_O_2_ feeding solution until dissection for respective timepoints. For the dissections mosquitoes were knocked down on ice, and placed into falcon tubes with 70% ethanol within the glove box, snap frozen at −80°C for at least 10min and transferred into 1x phosphate-buffered saline (PBS) before midguts (oocyst stage) or salivary glands (sporozoite stage) were dissected. Each experiment included at least three individual replicates and up to 5.

### Oocyst counts and size measurement

Midguts of *P. falciparum* infected mosquitoes were dissected 5 dpIBM and 10 dpIBM in PBS, incubated for 20min in nonidet P-40 detergent (Biozol, Germany) and stained in 0.1% mercurochrome (Sigma -Aldrich, Germany) for 40 min. After rinsing three times in PBS, midguts were left in PBS and imaged at 10x magnification. Oocysts were counted and measured in FIJI by using scaled images ^140^. Size measurement included only fully visible oocysts and burst oocysts were also excluded from the analysis.

### Longevity assays

*PY-R* (Tiefora) mosquitoes were kept in individual cups and provided with feeding solutions in cotton wool pads containing multiple L-arginine or hydrogen peroxide dilutions in 10% sucrose solution for determining a concentration that doesn’t change the longevity against control populations receiving 10% sucrose. Feeding solutions were refreshed every day and mortality was also scored daily.

### Salivary gland sporozoite quantification

#### gDNA extraction using DNAzol

Salivary glands were dissected 16 days post-infection and pools of 1-3 pairs were collected in 1.5 ml tubes containing 10µl of PBS. Salivary glands were homogenized with a pestle and remaining liquid washed off into the tube with another 10µl PBS to a total volume of 20µl. Samples were stored at −80°C until further processing. The gDNA was extracted with DNAzol. Briefly, Samples were incubated for 2 min at RT with 1 ml DNAzol and 10µl glycogen was added followed by 500µl ice-cold 100% ethanol. After incubation on ice for 10 min and centrifugation for 10 min at 4°C, supernatant was removed and the pellet washed twice with 1 ml 75% ethanol by inverting and centrifugation of the tubes for 5 min. Ethanol was removed, samples were allowed to air-dry and dissolved by incubation with 20µl molecular grade water at 65°C for 10 min.

#### Sporozoite quantification by qPCR

Known numbers of salivary gland sporozoites counted in a haemocytometer were serial diluted and used for generating a standard curve by isolating the gDNA with DNAzol ^141^. The qPCR was performed using specific oligonucleotide primer and a protocol for a large ribosomal subunit fragment (LSUE) encoded in in the mitochondrial genome of *P. berghei, P. yoelii*, and *P. falciparum* (www.plasmodb.org; Table S1) ^141^. For the qPCR analysis 4µl of the extracted gDNA were used in a total PCR mix volume of 20µl using BRYT Green® Dye (Promega, Germany) with the CFX96™ Real-Time System (Bio-Rad, Germany) that includes the CFX Maestro 1.1 software (Bio-Rad, Germany). The cycling protocol was 95 °C for 10 min initial denaturation followed by 40 cycles of: 95 °C, 15 s; 63.5 °C, 60 s.

### Immunofluorescence assay of P. falciparum ookinete invasion

Mosquitoes were snap frozen 26 hours post *P. falciparum* infection, midguts dissected and prefixed in 4% PFA in PBS for 1 min and opened longitudinally on ice-cold PBS to remove the blood bolus. Midguts were then fixed in 2%PFA in PBS at RT, washed in PBS and blocked for 1 hour in PBST buffer (1% BSA, 0.1% Triton X-100 in PBS) before incubation overnight at 4°C with polyclonal rabbit anti-nitrotyrosine primary antibody (Thermo Fisher scientific, Germany) 1:1000 in PBT. After washing on the next day, samples were incubated with Alexa Fluor™ 568 and mouse anti-Pfs25 antibody conjugated with Alexa Fluor 488 (Bousema lab, Radboud Medical Center, Netherlands), that stains ookinetes, zygotes and female gametes for 3-4 hours at RT. DNA was then stained with Hoechst 33342 (Thermo Fisher Scientific, Germany) for 30 min and midguts washed. Lastly, they were mounted with Vectashield® mounting medium (VWR, Cat. No.: VECTH-1000) on glass slides for confocal microscopy (Leica SP8) analysis.

### RNA interference using siRNA or dsRNA

#### Template generation for dsRNA

Specific primers were designed using NCBI primer blast to generate DNA templates with lengths between 300 – 600bp from *An. gambiae* s.l or *An. coluzzii* cDNA for double-stranded RNA (dsRNA) synthesis ^137^. All primers contained a T7 sequence (5’ – 3’: TAATACGACTCACTATAG) at the 5’ end of the strand and additional properties included a maximum of 3 consecutive nucleotides, GC content between 20 – 50% with self-dimer max ΔG: at 3’ end = −5, internal = −6. Designed primers were used for synthesis of *dsCAT* and dsGFP as non-target control (Table S1). PCR conditions were 98⁰C for 30 s, then 35 cycles of 98⁰C denaturation for 7 s, annealing at 55⁰C – 60.3°C for 10 s and elongation at 72⁰C for 60 s with a final extension for 5 min at 72⁰C. Specificity of products was confirmed by agarose gel electrophoresis and amplicons were purified using the QIAquick® PCR Purification following the manufacturer’s manual (QIAGEN, Germany). The dsRNAs were synthesised using MEGAscript™ T7 Transcription Kit (Thermo Fisher Scientific, Germany) by following the user manual with a 16-hour incubation at 37⁰C. Samples were purified using the MEGAclear™ Transcription Clean-Up Kit (Thermo Fisher Scientific, Germany) with a double elution step at 65⁰C for 10 minutes to achieve a 100 μl final elution volume. Quality and quantity of the products were measured on a NanoDrop Lite spectrophotometer (Thermo Scientific, Germany) and concentrated to 3 μg/μl in a vacuum centrifuge.

#### dsRNA injections

Mosquitoes were anaesthesised on CO_2_ gas and 69 nl of concentrated dsRNA was injected into the thorax between cuticle plates for RNA interference using pulled needles. To control for the injection-induced stress dsGFP was injected at the same concentration and volume. Efficiency of the dsRNAs was tested 72 hours post-injection by comparing transcript expression of the target genes using qPCR (see qPCR analysis section) with transcript expression compared to the dsGFP control. Survived Injected female mosquitoes were used for infection experiments with *P. falciparum*.

### 16S amplicon sequencing

Groups of 5 whole mosquitoes were pooled per treatment and DNA extracted by using the LIVAK protocol with a buffer of pH 7.5 containing 5M NaCl, 0.5M EDTA, 1M Tris pH 7.5 and 10% SDS added up to 100ml with ddH_2_O ^142^. Briefly, outer parts of mosquitoes were initially disinfected with 70% ethanol by vortexing and rinsed with sterile water. Dried mosquitoes were homogenized in 100µl 65°C warm LIVAK buffer and heated at 65°C for 30 minutes, followed by addition of 14µL 8M potassium acetate, mixing and 30 minutes incubation on ice. The supernatant was collected after centrifugation and 200µL pure ethanol (>99.9%) was added for precipitation of the DNA. Finally, the pellet was resuspended in 50µL ddH_2_O, concentration measured via Qubit and send to an external company (Biomarker Technologies (BMK) GmbH) for sequencing and provided 16S abundance tables analysis.

### Bulk RNAseq on whole P. falciparum infected mosquitoes

PY-R Tiefora received either L-Arg-, H_2_O_2_ feeding solution or 10% sucrose (Cntrl) continuously from emergence to adults onwards. At age 3-5 days all treatment groups were infected with *P. falciparum* gametocytes at 0.2% and snap frozen at −80°C exactly 21 hours post-infectious blood meal, around the time when ookinetes start invading the midgut epithelium. RNA was extracted using the PicoPureTM RNA-Isolation kit (see RNA extraction section), the RNA concentration and quality measured via Nanodrop lite spetrophotometer and Bioanalyzer 2100, and sequencing performed by external companies. Sequences were quality- and adapter-trimmed using Trimmomatic Version 0.38 ^143^ with default parameters and the quality was assessed using FastQC v0.11.9 (Andrews 2015). Resulting reads were mapped to the An. gambiae PEST v4 reference genome using HISAT2 v2.2.1 ^144^ and gene counts were obtained using featureCounts v 2.0.3 ^145^). Gene expression analysis was conducted with DESeq2 v4.3 ^146^ as previously described ^5^. Briefly, we filtered the genecounts data and kept only those transcripts with more than 10 reads in at least n-1 samples. Sample metadata was passed to DESeqDataSetFromMatrix, variance stabilised dispersions were calculated and PCAs and heatmaps were created on this dataset. Differential expression was calculated using DESeq with standard settings. Genes were considered significantly differentially expressed if p < 0.05 after correction and used for further analysis.

### Assessment of potential effects on malaria transmission using a compartmental model

#### Baseline model

In order to assess the potential effects of permethrin tarsal exposure or feeding L-arginine in the field on malaria transmission, a compartmental model approach was used. Three susceptible-exposed-infectious-recovered (SEIR) models were developed, each with a human and a mosquito population. The baseline model is intended to estimate the mosquito population dynamics and *Plasmodium falciparum* transmission under natural conditions in southern Burkina Faso. Since *Annopheles coluzzii* is a major vector there ^147^, the mosquito population was simulated with data from this species. The transmission of *P. falciparum* between the three populations and the developments are calculated by the dynamics between the compartments described using differential equations.

The total human population (N_h_) includes the subpopulation of susceptibles (S_h_), the subpopulation of exposed (E_h_), the subpopulation of infectious (I_m_) and the subpopulation of recovered (R_h_). The mosquito population consists of an aquatic population in which the mosquitoes develop and a total adult population (N_m_). The aquatic population consists of the developmental stages of the mosquito (eggs_m_), the mosquito larvae (lar_m_) and the mosquito pupae (pup_m_). The total adult mosquito population (N_m_) includes the subpopulation of susceptible (S_m_), the subpopulation of exposed (E_m_), the subpopulation of infectious (I_m_) and the subpopulation of resistant (R_m_).

As female mosquitoes are relevant for malaria transmission, only these are included in the model. The assumption is made that the sex ratio is equally distributed during oviposition. During a blood meal, contact between humans and mosquitoes can lead to transmission of *P. falciparum* ^148^. Two types of transmission are relevant for the model:

1. An infectious mosquito (I_m_) bites a susceptible human (S_h_) and transmits *P. falciparum* to the now exposed human (E_h_) with a transmission rate (beta_h_). After an incubation period (sigma_h_), an infected person becomes an infectious (I_h_) and recovers with a recovery rate (gamma_h_). After approx. 90 days, the recovered human (R_h_) becomes susceptible again (S_h_) ^149^.
2. An infectious human (I_h_) is bitten by a susceptible mosquito (S_m_). The mosquito is exposed to *Plasmodium* with a transmission rate (beta_m_). In this exposed mosquito E_m_, *Plasmodium* replicates after an extrinsic incubation period with an infection rate (sigma_m_) so that the mosquito becomes infectious (I_m_).

If the parasites are unable to replicate during the extrinsic incubation period, the exposed mosquito in the model is shifted to the subpopulation of resistant mosquitoes (Rm) with a resistance rate of (gamma_m_).

The development of the aquatic *An. coluzzii* stages and the corresponding transition from one compartment to the next compartment is calculated using development parameters such as oviposition, development time and survival time. When developing the model, climatic seasonality was taken into account on the basis of data from southern Burkina Faso in the form of a two-stage division in which the dry and rainy seasons alternate ^150^. In addition, density-dependent factors were included in the model, which have a negative effect on the longevity and reproduction of the population as the mosquito population grows, so that population growth is dynamically limited to a maximum population size. In addition, the following assumptions were made for the model: There is no vertical human malaria transmission. The human population is stable and there are neither births nor deaths. Humans are temporarily immune after surviving infection before becoming fully susceptible again. The model was simulated using R version 4.4.2 (R Core Team, 2024). The R package ‘deSolve’ version 1.40 was used to calculate the differential equations ^151^. The model is summarized in the following flow chart:

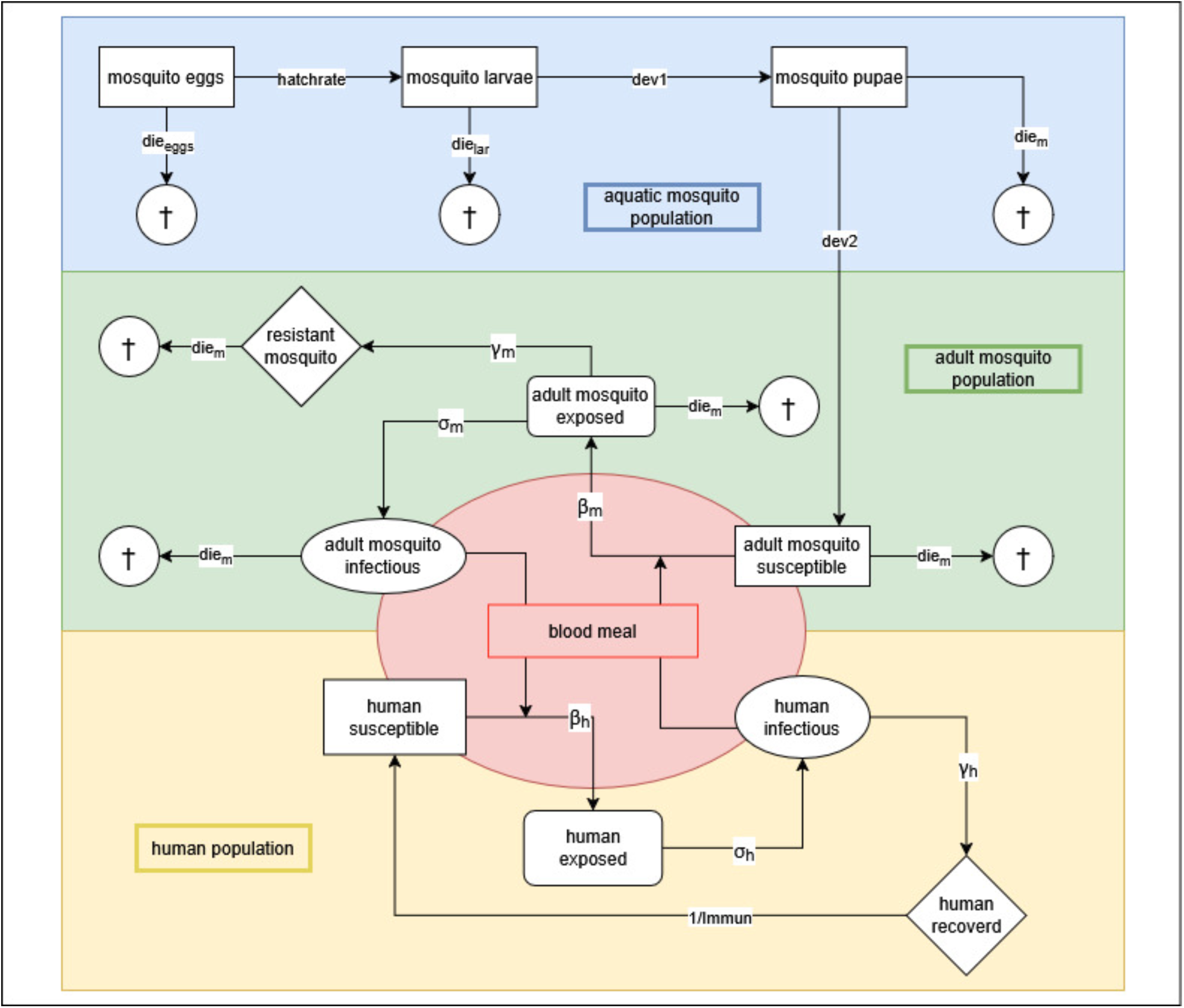

#### Treatment model

Based on the baseline model, two further models were programmed including oocyst prevalence data from permethrin and L-Arg treatment in PY-R mosquitoes individually. The treatment models were adjusted with the parameters determined in the laboratory for reproduction, survival and transmission of the mosquitoes from the different treatment groups. Comparison of the results of the treatment model with the baseline model shows the potential effect of permethrin or L-Arg interventions on malaria transmission under field conditions.

### Fecundity assays

Mosquitoes receiving an L-Arg, Glyc or Cntrl (10% sucrose) diet or 10% sucrose fed mosquitoes tarsal exposed to 0.01% permethrin in acetone or acetone-only coated glass dishes were provided an uninfectious blood meal at 3-5 days of age. The exposure was performed 3 hours before the blood-meal. 20-25 mosquitoes per group were kept in a cup and two days after the blood meal a vessel with wet filter paper was added for egg laying. The vessel was left in the cup for two days and egg numbers were counted 4 days after blood-feeding. For hatching rate, the number of hatched larvae was counted two days after egg laying and emergence rate was calculated by counting remaining dead larvae in trays after emergence to adults.

### Quantification and statistical analysis

Details of the statistical methods used in each experiment are included in the figure legends. Data was analyzed and graphs produced in GraphPad Prism 10 (GraphPad Software, La Jolla California USA). Normality of data was determined for all experiments using the D’Agostino-Pearson omnibus test with a cut-off value of p = 0.05 for significance. Generally, when data was not normally distributed it was checked for skew direction. When data was bimodal or irregular no transformation was attempted and when the data was right skewed transformation was applied via Log (base 10) and checked for normality again. In all cases where no transformation to normal distribution was possible for at least one of the compared groups non-parametric statistical tests were applied. A significance value of p = 0.05 was used as cut-off and Welch’s correction (t-test) or Dunnet’s multiple comparison test (ANOVA) applied as post-hoc tests for correction of the significance. Mann Whitney test was used for non-parametric individual comparisons. For all parametric and non-parametric t-test comparisons used, the significance threshold of p = 0.05 was additionally adapted by Bonferroni correction and displayed in figure legends. Proportional data (infection prevalence, hemocyte proportions, egg hatching rate, larval emergence rate, ookinete nitration, fragmentation) was compared using a chi-square test as n > 5 in all cases. The significance is presented in all graphs as p-value, while non-significant analyses, ns’ are only presented in proportion graphs or when multiple comparisons were done with more than one control group and otherwise left as empty space. Lines beneath p-value were only used when there were multiple control groups or multiple comparisons.

## Key resources table

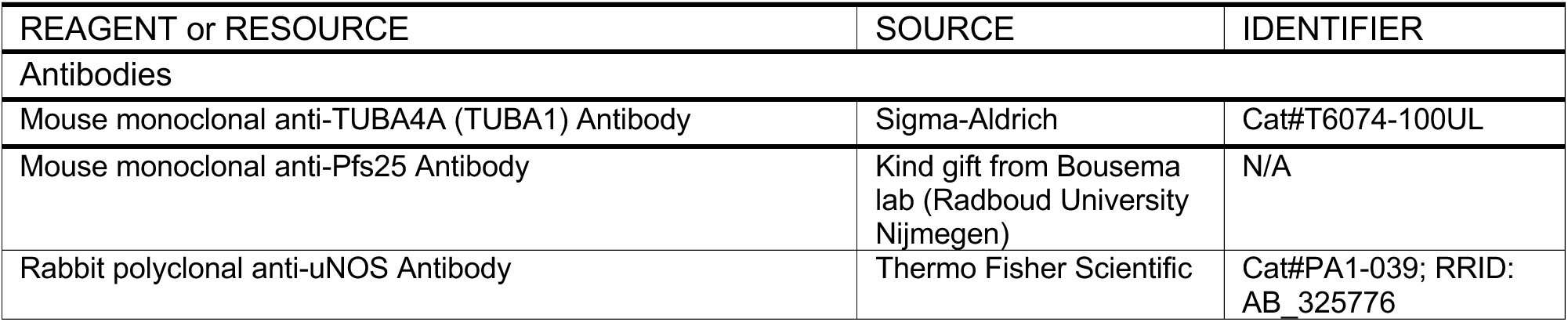
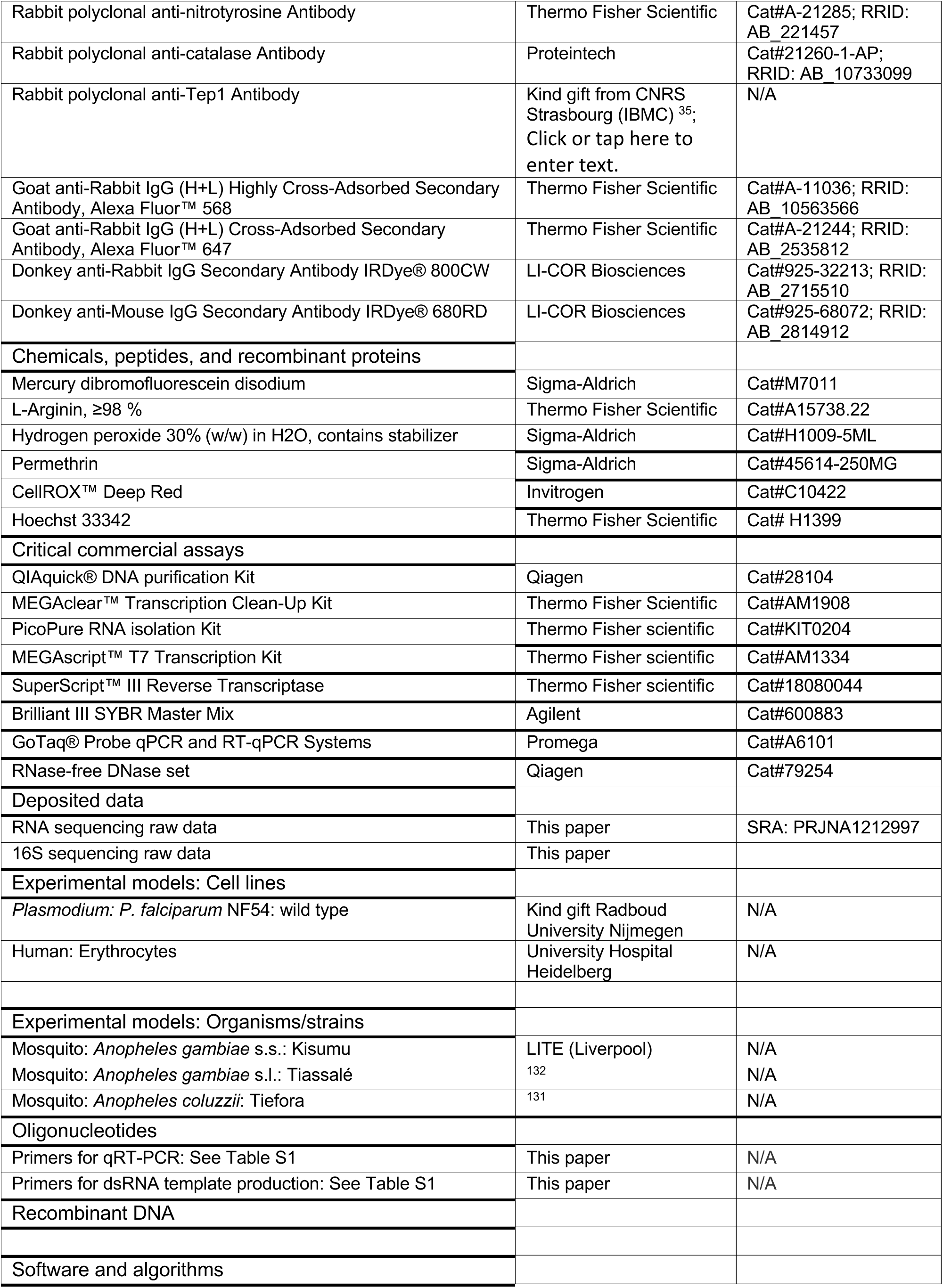
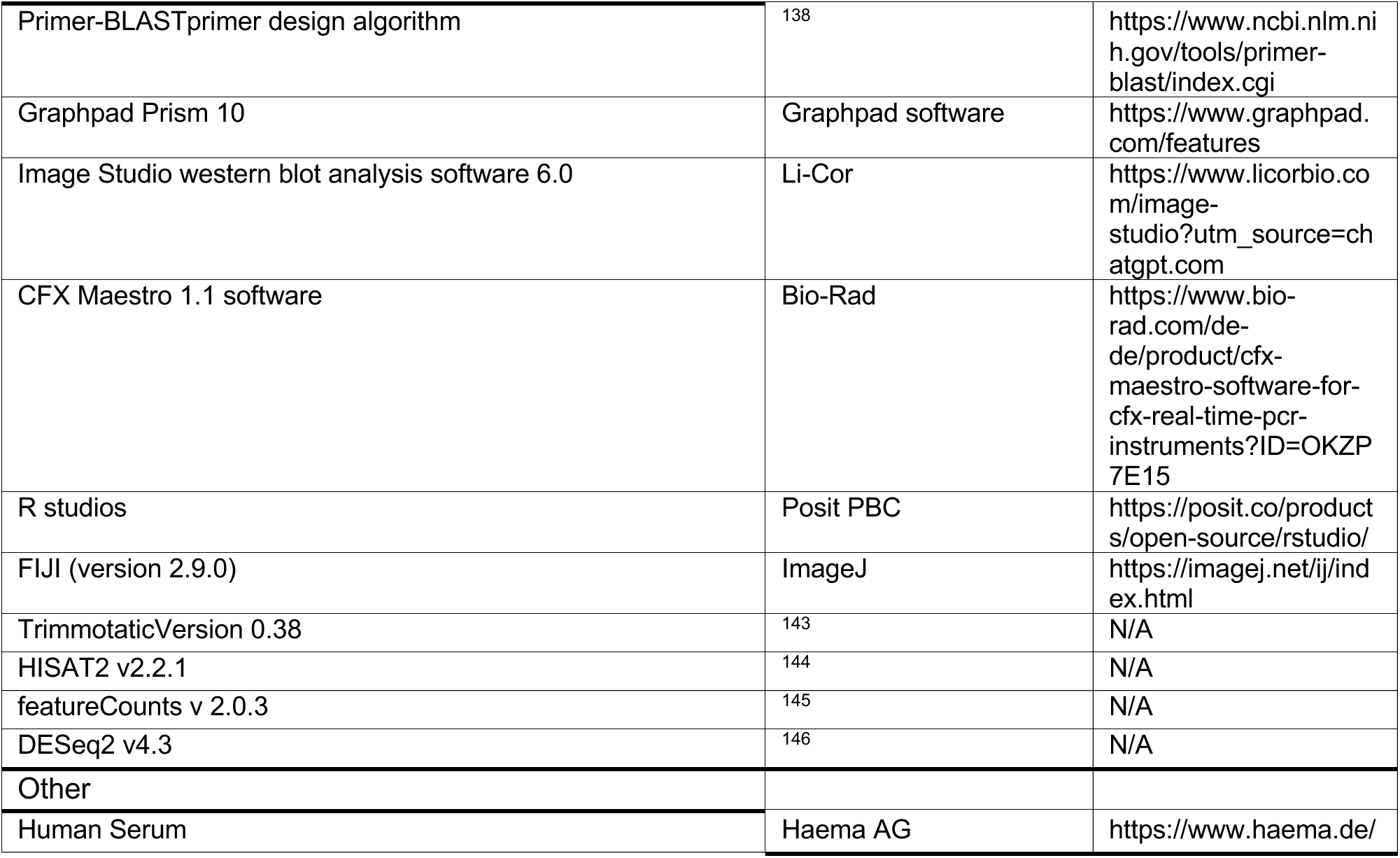

## Notes

### Competing Interest Statement

The authors have declared no competing interest.

